# Review and further developments in statistical corrections for Winner’s Curse in genetic association studies

**DOI:** 10.1101/2022.11.28.518299

**Authors:** Amanda Forde, Gibran Hemani, John Ferguson

## Abstract

Genome-wide association studies (GWAS) are commonly used to identify genomic variants that are associated with complex traits, and estimate the magnitude of this association for each variant. However, it has been widely observed that the association estimates of variants tend to be lower in a replication study than in the study that discovered those associations. A phenomenon known as *Winner’s Curse* is responsible for this upward bias present in association estimates of significant variants in the discovery study. We review existing *Winner’s Curse* correction methods which require only GWAS summary statistics in order to make adjustments. In addition, we propose modifications to improve existing methods and propose a novel approach which uses the parametric bootstrap. We evaluate and compare methods, first using a wide variety of simulated data sets and then, using real data sets for three different traits. The metric, estimated mean squared error (MSE) over significant SNPs, was primarily used for method assessment. Our results indicate that widely used conditional likelihood based methods tend to perform poorly. The other considered methods behave much more similarly, with our proposed bootstrap method demonstrating very competitive performance. To complement this review, we have developed an R package, ‘winnerscurse’ which can be used to implement these various *Winner’s Curse* adjustment methods to GWAS summary statistics.

**Author Summary:** A genome-wide association study is designed to analyse many common genetic variants in thousands of samples and identify which variants are associated with a trait of interest. It provides estimates of association strength for each variant and variants are classified as associated if their test statistics obtained in the study pass a chosen significance threshold. However, due to a phenomenon known as *Winner’s Curse,* the association estimates of these significant variants tend to be upward biased and greater in magnitude than their true values. Naturally, this bias has adverse consequences for downstream statistical techniques which use these estimates. In this paper, we look at current methods which have been designed to combat *Winner’s Curse* and propose modifications to these methods in order to improve performance. Using a wide variety of simulated data sets as well as real data, we perform a thorough evaluation of these methods. We use a metric which allows us to identify which methods, on average, produce adjusted estimates for significant variants that are closest to the true values. To accompany our work, we have created an R package, ‘winnerscurse’, which allows users to easily apply *Winner’s Curse* correction methods to their data sets.

## Introduction

It has been observed that in general, the effect size of a variant or single nucleotide polymorphism (SNP) tends to be lower in a replication study than in the genome-wide association study (GWAS) that discovered the SNP-trait association. This observation is due to the phenomenon known as *Winner’s Curse.* In the context of a single discovery GWAS, the term *Winner’s Curse* describes how the estimates of association strength for SNPs that have been deemed most significant are very likely to be exaggerated compared with their true underlying values. These estimated effect sizes can take the form of log odds ratios (log-OR) resulting from a logistic regression for a binary outcome, e.g. disease status, or regression coefficients (beta) derived from a linear regression for a quantitative trait.

Dudbridge & Newcombe (1) detail two sources of *Winner’s Curse* in GWASs, namely ranking bias and selection bias. Ranking bias stems from ranking many SNPs, often close to a million or more, by some measure of effect size or statistical significance. In practice, *p*-values are generally used. It is then expected that the bias will be greatest for those variants which have been ranked highly. Selection bias describes how the use of a stringent threshold, such as 5 × 10^-8^, can result in overestimated effect sizes for SNPs that exceed this threshold.

*Winner’s Curse* bias can have many practical consequences, especially with respect to techniques which are reliant on SNP-trait association estimates obtained from GWASs. One such example is Mendelian randomization (MR), a statistical framework which uses genetic variants as instrumental variables to estimate the magnitude of the casual effect of an exposure on an outcome. In the case of two-sample MR, if the same GWAS is used to identify instrument SNPs and estimate their effects relative to the exposure, *Winner’s Curse* will result in the over-estimation of these SNP-exposure associations. This bias will then propagate into the causal estimate, resulting in a deflation of this estimate. On the other hand, if instrument SNPs are discovered in the same GWAS as that used to estimate the SNP-outcome associations, the causal estimate will be inflated (2). In addition, *Winner’s Curse* has been shown to greatly increase the magnitude of weak instrument bias in these MR analyses (3). Another implication of *Winner’s Curse* bias is in the use of polygenic risk scores which employs GWAS results for prediction purposes. Enlarged association estimates of significant variants used in creating the polygenic score can lead to reduced accuracy in out-of-sample prediction (4).

In this paper, we review existing *Winner’s Curse* correction methods and explore possible modifications that could be made in order to improve these methods. However, eliminating this bias induced by *Winner’s Curse* is known to be a difficult task. Several bias reduction approaches have been proposed in recent years, with one of the earliest being the Conditional Likelihood method suggested by Ghosh et al. (5). This method makes an adjustment to the association estimate of each SNP which has been deemed significant, i.e. those with *p*-values less than the specified genome-wide significance threshold. In contrast to this approach in which the correction is performed to each SNP separately, independently of estimated associations of other SNPs, alternative methods have been suggested which involve the use of all SNPs, including those which do not pass the threshold, in order to produce bias-reduced estimated effect sizes. The empirical Bayes method described by Ferguson et al. (6) determines a suitable correction for each SNP by using the collective distribution of all effect sizes. Bigdeli et al. (7) suggested the use of FDR Inverse Quantile Transformation (FIQT), while Faye et al. (8) proposed a bootstrap shrinkage estimator with application to the GWAS setting. As this bootstrap approach requires individual-level data, we propose an alternative form of this method which uses bootstrapping with summary statistics to make corrections.

The focus in this paper is on methods which attempt to reduce the effect of the bias induced by *Winner’s Curse* using only GWAS summary statistics, not individual-level data, the reason being that approaches based on summary data tend to be more computationally efficient, in terms of run time and memory efficiency. Furthermore, GWAS summary statistics are much more accessible and are more widely used in epidemiological techniques such as MR. In addition, there exist methods which use both a discovery and a replication GWAS in order to make suitable corrections to estimated effect sizes of significant SNPs. Examples include the UMVCUE of Bowden and Dudbridge (9) and an additional conditional likelihood method, Zhong and Prentice (10). That said, the concentration of our work detailed here is on techniques which have been designed for use when a replication sample is unavailable.

As mentioned above, we have made amendments to existing *Winner’s Curse* correction methods to address certain weaknesses. In particular, we investigated modifications that could be made to the empirical Bayes method in order to ensure that it makes better adjustments to association estimates. Following this review of correction methods, a rigorous evaluation and comparison of these methods was performed. This assessment took place by means of a simulation study as well as engagement with three real data sets. Simulations allowed us to compare methods easily over a wide range of different possible genetic architectures. We then used UK Biobank (UKBB) body mass index (BMI), type 2 diabetes (T2D) and height data sets to see how these techniques would perform in more realistic settings in which a large degree of linkage disequilibrium (LD) exists. In both instances, assessment of methods was predominantly based on the computation of estimated mean squared error (MSE) over significant SNPs. A notable challenge that was encountered at the start of the work discussed in this paper was the lack of available software to implement these various correction methods. Therefore, to complement this review, we have developed an R package, namely ‘winnerscurse’ (https://github.com/amandaforde/winnerscurse), which can be used to apply a number of *Winner’s Curse* adjustment methods to GWAS summary statistics. Techniques which require a replication GWAS are also included in this package.

## Materials and methods

Throughout this paper, we let 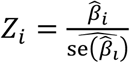 and 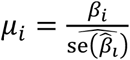 with the assumption that

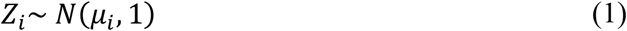

asymptotically, in which *β_i_* denotes the true effect size of SNP *i* for 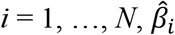 its estimated effect size, with respect to a trait of interest, and 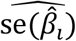 its estimated standard error. *N* represents the total number of SNPs in the discovery GWAS. Depending on the type of phenotype, be it a disease or a quantitative trait, this estimated effect size can represent a log odds ratio or a regression coefficient attained from a linear regression, respectively.

### Conditional Likelihood

As mentioned, the conditional likelihood method of Ghosh et al. (5) notably differs in its approach at making *Winner’s Curse* corrections from the other methods evaluated in this paper. Adjustments are made to the estimated effect sizes of only those SNPs which satisfy |*z*| > *c*, where *c* is the value corresponding to the pre-specified significance threshold. The reduction in estimated effect size for each significant SNP is imposed independently of other SNPs and is directly determined by the value of *c*. Recognizing that a SNP has been deemed significant, the corresponding conditional likelihood is given by:

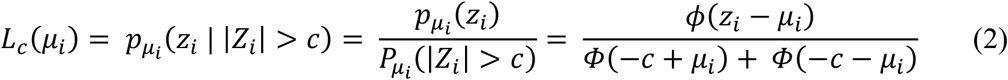

in which 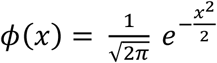, the probability density function of the standard Gaussian distribution and 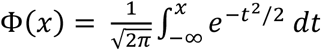, the corresponding cumulative distribution function (cdf). In general, *c* takes the form of 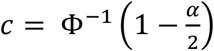, with *α* being a threshold to which a Bonferroni correction has been applied in order to control for family-wise error rate, e.g. 5 × 10^-8^.

Using this conditional likelihood, three estimators of *μ_i_*, or equivalently of *β_i_* as any estimator for *μ_i_* can be used to produce an estimator for *β_i_* by simply multiplying it by 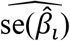, were proposed. The first, 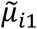 is the obvious conditional maximum likelihood estimator:

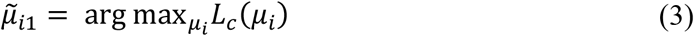

while the second, 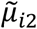 is defined as

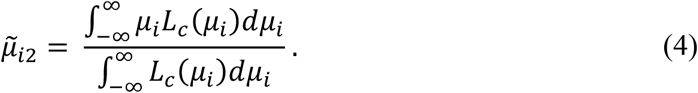

This is the mean of the random variable that follows the distribution *L_c_*(*μ_i_*), normalized to ensure a proper density. However, it was observed that for instances in which the true effect size, *β_i_* is close to that of a null effect, the estimator 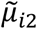 has greater mean squared error than 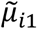 but for true effect sizes further from zero, 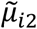 performs better. Therefore, the use of 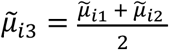, which can combine the strengths of these two estimators in order to curtail *Winner’s Curse* bias for significant SNPs more accurately, was suggested.

### Empirical Bayes

Motivated by Efron’s empirical Bayes implementation of Tweedie’s formula to correct for selection bias (11), the empirical Bayes method detailed by Ferguson et al. (6) focuses on the importance of sharing information between SNPs in order to make adjustments, through the exploitation of the empirical distribution of all effect sizes. This is a notably different approach to that of the previously discussed conditional likelihood method, which when making a correction to the estimated effect size of a particular SNP essentially fails to acknowledge the existence of any other SNPs.

Under the normal sampling assumption described by Eq (16), Tweedie’s formula describes the relationship between the posterior mean, *E*(*μ*|*z*), and the marginal density function, *p*(*z*), as

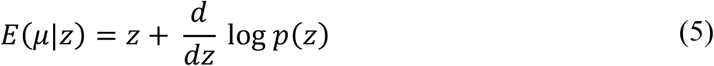

Amazingly, provided one can estimate *p*(*z*), Tweedie’s formula facilitates estimation of the posterior mean in complete absence of knowledge of the prior distribution, *p*(*μ*), which in this instance is the true distribution of standardized effect sizes across the genome. Thus, the estimator of *μ_i_*, proposed by this method takes the form of

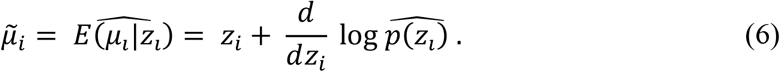

Estimation of log *p*(*z*) occurs upon application of the following steps. First, partitions of the interval [*z*_1_, *z_N_*] of identical width are formed, in which the *z*-statistics have been arranged in ascending order. The number of *z*-statistics which fall inside each partition are noted and regressed against a set of natural cubic spline basis functions with knots located at the midpoint of each partition, using a Poisson generalized linear model. Ferguson et al. (6) suggest choosing the number of basis functions so that the Bayesian Information Criterion (BIC) is minimized for the model. The fitted regression function at *z* is then used to obtain the estimate for log *p*(*z*) and subsequently, 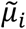 for *i* = 1, …,*N*, by means of numerical differentiation.

Ferguson et al. (6) show that if the true marginal density, *p*(*z*), could be used here, then the empirical Bayes estimator would perform optimally at minimizing the mean squared error (MSE) over all SNPs. However, since it is only an estimate of *p*(*z*) that can be obtained, this optimal behaviour is not guaranteed. This is especially a concern in the extreme tails of the distribution where the z-statistics of the most significant SNPs lie as it is more difficult to accurately estimate *p*(*z*) in these regions. Ferguson et al. (6) considered an ad hoc strategy to assist in overcoming this issue. The suggested approach involves the combination of this estimator with the conditional likelihood estimator, in a manner which is determined by the estimators’ respective lengths of 95% confidence and credible intervals. Here, we instead investigated 5 alternative modifications to the original empirical Bayes method described above in order to better stabilize the tail of the estimated marginal density, 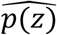 and its derivative, particularly in the context of strong LD that is observed in high density genotyping arrays. These variations avoid the unappealing combination of two appreciably different estimators, empirical Bayes and conditional likelihood. The explored modifications were:

- Altering the minimum-BIC estimated spline function to be log-linear beyond the 10^th^ largest negative and the 10^th^ largest positive *z*-statistics
- Limiting the number of knots in the spline, in particular using 7 degrees of freedom as originally suggested by Efron (11)
- Utilizing smoothing splines, rather than natural splines, through the gam function in the R package mgcv (12), to avoid specifying knot positions, assuming a poisson distribution for the partition counts
- As above, but this time using a more realistic negative binomial distribution for these counts
- Employing splines with additional shape constraints, through the scam function in the R package scam (13), to enforce monotonicity of the estimated density function, 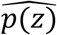

More information about these modifications and their rationale is given in the supplementary material.

### FDR Inverse Quantile Transformation

FDR Inverse Quantile Transformation (FIQT), as proposed by Bigdeli et al. (7), employs a straightforward two-step procedure in order to produce less biased association estimates. First, a FDR (false discovery rate) multiple testing adjustment is applied to the *p*- values of all SNPs, giving FDR adjusted *p*-values *p_i_*\*, *i* = 1, …,*N*. Following this, these adjusted *p*-values are transformed back to the z-statistic scale by means of an inverse Gaussian cumulative distribution function (cdf) and for each SNP, it is ensured that this new *z*-statistic, 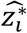, has the same sign as its original effect size. Mathematically, 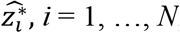, can be described as

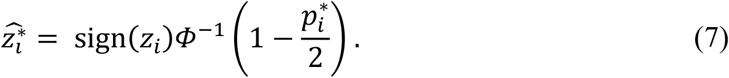

For SNP *i*, its new estimated effect size is simply calculated as 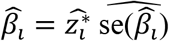.

The rationale that led to the use of this method is based on the analogy between performing multiple testing adjustments to *p*-values and reducing *Winner’s Curse* bias in estimated SNP effect sizes, in which these effect sizes are in the form of *z*-statistics. In the attempt to correct for *Winner’s Curse,* a shrinkage towards the null effect of zero is generally incurred by the *z*-statistics while the application of a multiple testing adjustment to *p*-values sees the growth of the *p*-values towards one, the null value.

This multiple testing adjustment is imposed through the implementation of the R function p.adjust. This is followed by the use of the R function qnorm for the purpose of back-transformation. However, near zero *p*-values can prove problematic when evaluating qnorm and thus, a restraint is incorporated in FIQT which results in the association estimates of SNPs with very large *z*-statistics, e.g. greater than 37, failing to be adjusted.

### Bootstrap

Inspired by the bootstrap resampling method detailed in Sun et al. (14), we have established a similar approach which can be easily applied to published sets of GWAS summary statistics without requiring original individual-level data. In addition, a second advantage of our new method is a considerable improvement in computational efficiency over the method described in Sun et al. (14).

This procedure begins with arranging all *N* SNPs according to their original *z*- statistics, 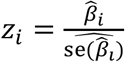, in descending order, that is a labelling of SNPs is assumed such that *z*_1_ > *z*_2_ > … > *z_N_*. A randomized estimate of the extent of ranking bias for the *k*^th^ largest *z*-statistic is calculated by means of the parametric bootstrap as follows:

1. A value 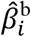 is simulated for SNP *i, i* = 1, … *N*, independently, from a Gaussian distribution with mean 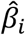 and standard deviation 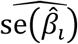, i.e.

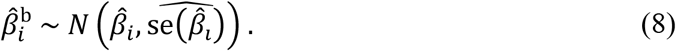
2. Upon obtaining 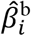 for *i* = 1,…, *N*, the 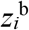-statistic of SNP *i* is defined as

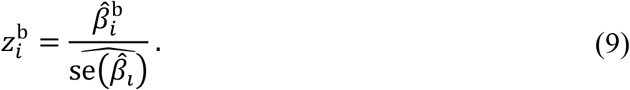 We define *A*(*k*) as the index corresponding to the *k*^th^ largest entry in the vector:
3. Then, the estimated bias of SNP *k,* the SNP with the *k*^th^ largest original *z*-statistic, takes the following form:

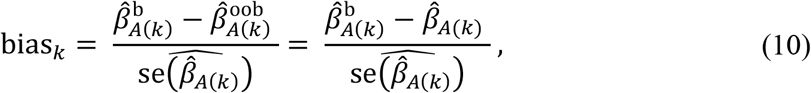

in which 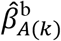 is the bootstrap value of the SNP ranked in position *k* in the ordering of 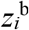-statistics, 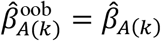 is that same SNP’s original *β* estimate and 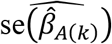 its standard error.

In the next step of the process, a cubic smoothing spline is fitted to the data in which the *z*- statistics are considered as the inputs and bias_*k*_, their corresponding outputs. The predicted values from this model fitting provides new estimates for the bias correction, bias*k*\* for each SNP. This additional stage in which bias_*k*_* is obtained reduces the need for more than one bootstrap iteration for each SNP in order to ensure competitive performance of the method. This results in a faster approach with increased accuracy. Finally, the new estimate for the true effect size of SNP *k*, the SNP with the *k*^th^ largest original *z*-statistic, is defined as: 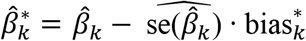.

In addition to those mentioned previously, there are several notable differences between our algorithm described above and the method proposed by Sun et al. (14). Firstly, it is the parametric bootstrap that is used here to estimate the magnitude of bias for each SNP as opposed to the more common nonparametric bootstrap which requires individual-level data. Our method draws only one bootstrap resample, i.e. only one bootstrap value 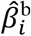 is simulated for SNP *i, i* = 1, …,*N*. It also includes an extra step which involves the use of a smoothing spline. In contrast, Sun et al. (14) express the need for a number of bootstrap samples, e.g. at least 100, in their approach. Furthermore, our algorithm based on the parametric bootstrap only corrects for ranking bias, and not threshold-selection bias.

### Simulation study

The simulation study followed a factorial design in which GWAS summary statistics were simulated for a quantitative trait under 8 different genetic architectures, described by combinations of three parameters, namely sample size *n*, heritability *h*^2^, polygenicity (proportion of effect SNPs) *π*. The following the values chosen for these parameters:

- sample size *n* ∈ {30000,300000}
- heritability h^2^ ∈ {0.3,0.8}
- polygenicity *π* ∈ {0.01,0.001}

Assuming a selection coefficient equal to zero and a normal distribution of effect sizes, for a fixed array of *N* = 1,000,000 SNPs, our strategy entailed imposing a simple correlation structure on the SNPs in order to imitate the presence of linkage disequilibrium (LD) in real data. It was assumed that the same correlation structure exists in independent blocks of 100 SNPs. Thus, for each block of 100 SNPs, the estimated effect sizes, 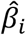 were simulated using:

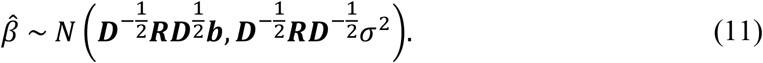

Here, ***b*** is a vector containing the true SNP-trait effect sizes which have been scaled to ensure that the phenotype has variance 1, i.e. *σ*^2^ = 1. The matrix ***D*** is a diagonal 100 × 100 matrix, in which *d_i_*=*n*· 2· maf_*i*_(1 – maf_*i*_) and maf*i* is the minor allele frequency of SNP *i*, while ***R*** is a simple 100 × 100 matrix of inter-genotype correlations, with 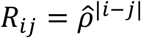 and 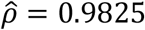. The reasoning for the selection of this value for 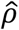 and why it was considered suitable, as well as other details regarding this simulation, are described in the supplementary material. For each SNP, values for 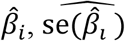 and 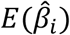 were produced with 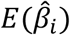 obtained using 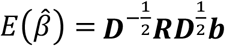. For each of these 8 different genetic architectures, 100 sets of summary statistics were simulated.

The *Winner’s Curse* correction methods detailed in ‘Materials and methods’ were applied to each data set using the R package ‘winnerscurse’, producing adjusted estimated effect sizes, 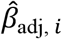, for each SNP *i, i* = 1, …,*N*. The performance of these methods were investigated at two different significance thresholds, namely *α*_1_ = 5 × 10^-8^ and *α*_2_ = 5 × 10^-4^, with a stronger focus given to the more commonly used genome-wide significance threshold of *α*_1_ = 5 × 10^-8^. In order to assess each method’s ability at providing less biased SNP-trait association estimates, both the estimated change in mean squared error (MSE) and estimated change in root mean squared error (RMSE) of significant SNPs due to method implementation were computed for each data set and method. For simplicity, let *i* = 1, …, *N*_sig_ represent indexes for the significant SNPs in a particular simulated set of summary statistics, i.e. *N*_sig_ is the number of SNPs which satisfy |*z_i_*| > *c* with 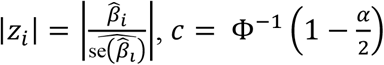 and *α* ∈ {*α*_1_, *α*_2_}. Then, the estimated change in MSE of significant SNPs may be defined as:

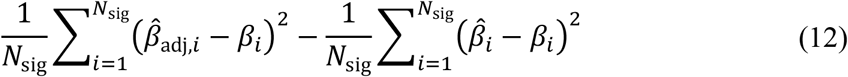

while the estimated change in RMSE is defined similarly as:

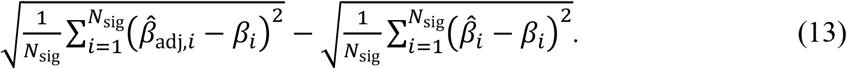

The change in MSE and RMSE for each method was calculated for only those data sets in which at least one significant SNP was detected. In addition to these two metrics, the relative change in MSE, which is equal to the change in MSE divided by the naïve MSE, was computed in a similar manner. For a given correction method, this value provides the percentage improvement in MSE due to applying that method to the set of summary statistics.

In addition to the above simulation set-up, GWAS summary statistics were simulated and methods evaluated under the assumption that SNPs were independent. In this instance, the study was extended to 24 genetic architectures in which the selection coefficient *S* took values −1 and 1 as well as 0. This simulation process which incorporates an independence assumption was repeated in a similar fashion for a binary trait with a normal distribution of effect sizes. Furthermore, a quantitative phenotype with a bimodal effect size distribution as well as one with a skewed distribution were also considered. In order to reduce computation time, only 50 sets of summary statistics were simulated for each combination of the four parameters for these three additional situations.

### Empirical analysis

In order to compare the performance of these *Winner’s Curse* correction methods with respect to real data, three different UK Biobank data sets were used, namely body mass index (BMI), height and type 2 diabetes (T2D). As with real data, the true effect size of each SNP is unknown and so it is more difficult to assess how much each method reduces the bias induced by *Winner’s Curse.* To overcome this challenge, each original large data set was randomly split in two, leaving between 166,172 and 166,687 individuals in each of the six smaller data sets. This provided the ability to execute two independent GWASs of similar sample size for each trait in which one GWAS was designated as the discovery GWAS and the other the replication GWAS. The unbiased replication GWAS association estimates can then be used as proxies for the true effect sizes of the SNPs found to be significant in the discovery GWAS. PLINK 2.0 (15) was used to perform quality control as well as each of the statistical analyses.

The required quality control steps which took place beforehand included the removal of related individuals. Samples which had been identified as outliers with respect to heterozygosity and missingness together with samples with discordant sex information and those suffering from chromosomal aneuploidy were also discarded. Furthermore, non-European samples which were identified by principal component analysis (PCA) using 1000 Genomes data were removed. With respect to variants, only those with an information score greater than 0.8, a minor allele frequency greater than 0.01, a genotyping rate of at least 98% and those that passed the Hardy-Weinberg test at the specified significance threshold of 1 × 10^-8^ were included.

The methods of interest were applied to the summary statistics of each discovery GWAS using the R package ‘winnerscurse’. Evaluation took place by computing the estimated MSE of *N*_sig_ significant SNPs in that GWAS, defined as:

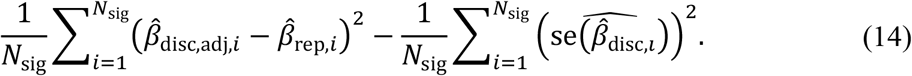

For each of the three traits, it was possible to evaluate the performance of methods twice as in each case, the original roles of the two independent data sets, i.e. discovery and replication, could be switched and re-evaluation of methods could then take place with respect to the SNPs that were deemed significant in this new discovery GWAS.

## Results

### Simulation study

#### When is winner’s curse bias most prominent?

A simulation study in which a simple correlation structure was imposed on the set of *N* = 1,000,000 SNPs was first executed, as described in ‘Materials and methods’. Before application of the *Winner’s Curse* correction methods to the sets of summary statistics, an attempt to gain an insight into the simulation scenarios in which *Winner’s Curse* bias is most prominent was made. This was done by computing the average number of significant SNPs, the average naïve MSE of significant SNPs and the average proportion of significant SNPs that had significantly overestimated effect sizes in each setting with respect to two significance thresholds, 5 × 10^-8^ and 5 × 10^-4^. A SNP is defined as being significantly overestimated or as having a significantly more extreme effect size estimate if it satisfies the condition:

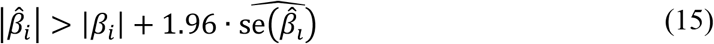

Thus, the proportion of significant SNPs that are significantly overestimated is considered to be representative of the proportion of significant SNPs with effect size estimates that greatly suffer from *Winner’s Curse* bias. As detailed in S1 Table, it was clear that as sample size was increased from 30,000 to 300,000, this proportion of significant SNPs decreased. The other two parameters which played key roles in defining the various simulated genetic architectures were heritability and polygenicity. It was observed that the proportion of significantly overestimated significant SNPs decreased when heritability was increased from 0.3 to 0.8, but when the value representing trait polygenicity was increased from 0.001 to 0.01, this proportion decreased.

In fact, it was also noted that as the number of significant SNPs increased, both the MSE of significant SNPs and the proportion of these that were significantly overestimated decreased. This can be clearly seen in S1 Fig. This indicates that as the number of samples in a study and as the number of SNPs passing the significance threshold increases, bias induced by *Winner’s Curse* will be less of an issue among significant SNPs. In terms of genetic architecture characteristics, these results suggest that the presence of *Winner’s Curse* bias in the estimated effect sizes of significant SNPs should be of a greater concern when investigating traits with lower heritability or traits which have a larger proportion of effect SNPs.

#### Evaluation of performance at *p* < 5 × 10^-8^

With respect to the evaluation of methods, we focus on the results of computing the quantity ‘change in RMSE over all significant SNPs due to method implementation’ for each method, with obtaining a negative value being desirable. These results are provided in S2 and S5 Tables. At a threshold of 5 × 10^-8^, several observations were notable. Firstly, for scenarios in which sample size has been designated the greater value of 300,000, the effect of applying the methods is on a much smaller scale to those scenarios with sample sizes of 30,000. This is evident from the large difference in the values on the y-axis between plots (A) and (B) of Fig 1. This observation ties in with the fact that the magnitude of *Winner’s Curse* bias is greater at *n* = 30,000. At this 5 × 10^-8^ significance threshold, the conditional likelihood methods are seen to perform poorly, especially when sample sizes are increased to 300,000. In most instances, these methods provide worse association estimates than the naïve approach, often increasing the RMSE. The reason for this observation is over-correction of estimated effect sizes, especially those that lie close to the significance threshold.

**Fig 1.**
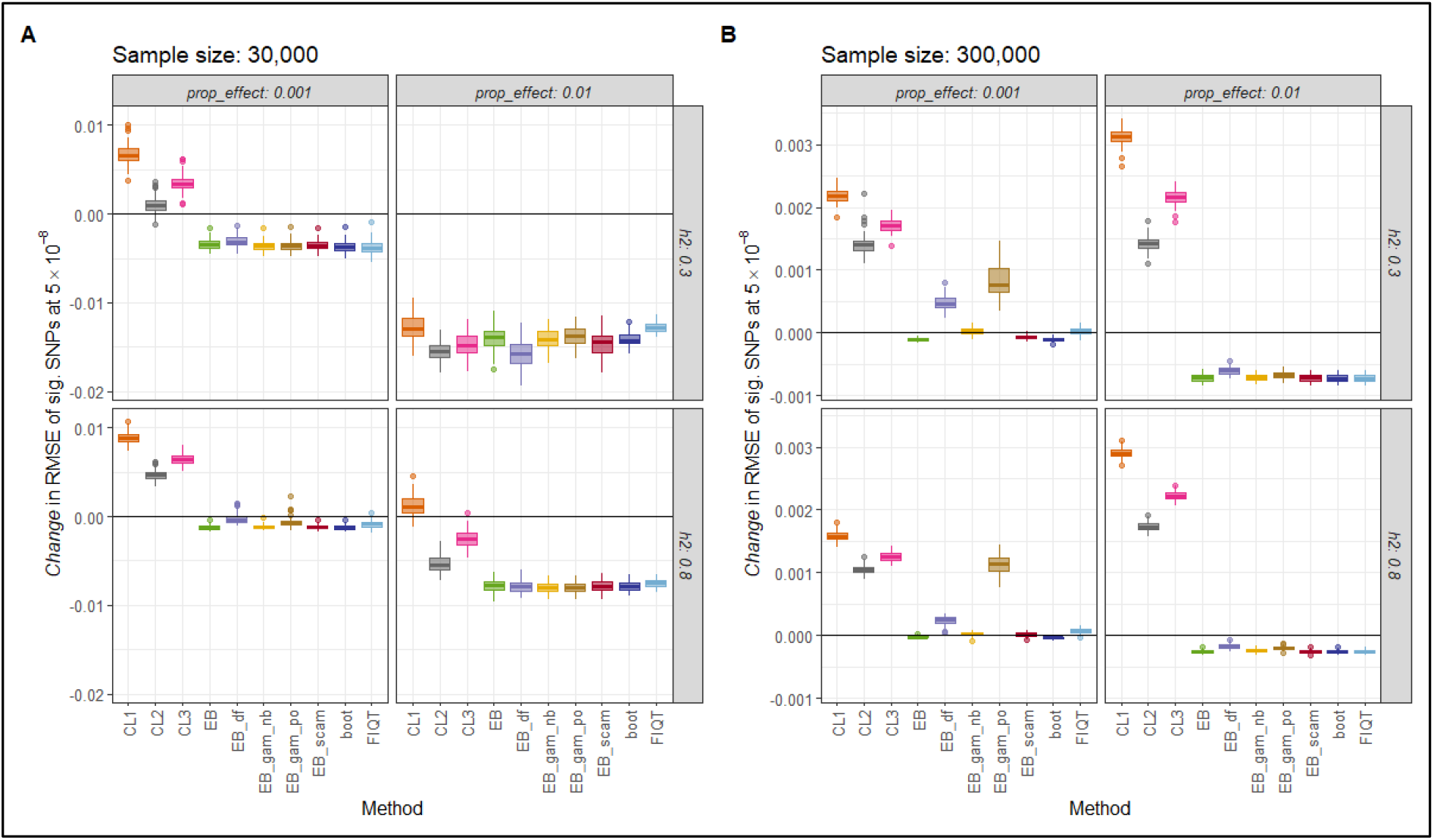
Estimated change in RMSE of significant SNPs at threshold 5 × 10^-8^ for each method and simulation setting, with a simple correlation structure imposed on the set of SNPs. The estimated change in RMSE of significant SNPs at a threshold of 5 × 10^-8^ (y-axis), as defined by Eq (13), is plotted for each correction method (x-axis), for each of the eight simulation settings. This figure corresponds to simulation settings in which a simple correlation structure has been imposed on the set of SNPs. Two four-panel plots, (A) and (B), are shown in the figure, in which (A) contains results related to settings with a sample size of 30,000 and (B) contains results for sample sizes of 300,000. The rows of these multi-panel plots represent heritability and the columns represent the proportion of effect SNPs. Each panel contains a boxplot for each correction method. As 100 sets of summary statistics were simulated for each simulation setting, an individual boxplot displays the distribution of estimated change in RMSE values obtained across the 100 sets with respect to a particular method and setting. The solid black line in each panel, representing no change in RMSE, is included in order to highlight which methods consistently provide negative values for the estimated change in RMSE. These methods are considered to be the best performing *Winner’s Curse* correction methods.

The novel bootstrap method is one of the most consistent methods at providing less biased SNP-trait association estimates at this threshold. In all situations depicted, it has one of the largest negative values for the change in RMSE of significant SNPs and on average, improves the MSE by 26% across the 8 settings, as shown in S4 Table. FIQT tends to perform in a somewhat similar manner to this bootstrap method. With respect to the empirical Bayes method and its variations, for sample sizes of 300,000, the best performing versions were the original empirical Bayes method, ‘EB-gam-nb’ and ‘EB-scam’. Note that the original empirical Bayes method is the form most similar to that proposed in Ferguson et al. (6) which also includes a restriction on the tails of the distribution of *z*-statistics. With the lower sample size, all variations perform very similarly. However, the empirical Bayes variants ‘EB-df’ and ‘EB-gam-po’ performed less well overall. In the case of ‘EB-gam-po’, this might be partly due to convergence problems that sometimes occurred in obtaining the poisson regression fit. In general, we advise caution in utilizing the empirical Bayes results in the context of convergence warnings from R.

#### Evaluation of performance at *p* < 5 × 10^-4^

A lower threshold of 5 × 10^-4^ was also investigated. Less emphasis is placed on the results obtained at this threshold as it is possible that many false positives are detected here, i.e. many SNPs that in fact have a true effect size of zero pass the significance threshold, although lower thresholds may be useful for the construction of polygenic risk scores. Therefore, as these *Winner’s Curse* correction methods are all considered to be shrinkage methods, improvements in the RMSE over all significant SNPs would be expected. However, as can be seen in S3 Fig, positive values are witnessed at this threshold when sample sizes are large. However, these positive values most often occur for the conditional likelihood methods which seem to be the worst performers overall. It seems here that the most consistent and best performing methods are the bootstrap method, the original empirical Bayes method and the empirical Bayes method which uses shape constrained additive models (SCAMs). These methods all reduce the MSE of significant SNPs at this threshold by an average of at least 32%, as shown in S7 Table.

#### Additional simulations in absence of Linkage Disequilibrium

In order to demonstrate potential performance of the *Winner’s Curse* methods in the context of SNP-trait associations from genome-wide arrays with lower SNP density or LD-pruned datasets, we also examined the less complex situation in which SNPs are independent. The results of these extra simulations are shown in S4-S11 Figs and described in depth in the supplementary material. In this setting, with a normal effect size distribution, the most consistent methods in terms of reducing the RMSE of significant SNPs were the original empirical Bayes method and ‘EB-gam-nb’, the variation of the empirical Bayes method which employs smoothing splines and assumes a negative binomial count distribution. Just as was observed in the simulations with linkage disequilibrium, the conditional likelihood methods perform poorly and often result in an increase in the evaluation metric in comparison to the naïve approach, while the proposed bootstrap method continued to exhibit competitive performance.

### Empirical analysis

The results of an initial exploration of the six UK Biobank sets of summary statistics are detailed in Table 1. From trait to trait, there is a large difference in the number of SNPs with *p*-values lower than the genome-wide significance threshold of 5 × 10^-8^. Values for the proportion of these SNPs with significantly overestimated effect sizes in each discovery GWAS are included. A comparison of BMI and height GWASs at the 5 × 10^-8^ threshold tends to indicate that as the number of significant SNPs increases, the proportion that are significantly overestimated decreases. This trend is even more apparent at the larger threshold of 5 × 10^-4^, and as stated above, was also clearly observed in the simulated data.

**Table 1.** The number of significant SNPs at two significance thresholds, 5 × 10^-8^ and 5 × 10^-4^, with proportions that indicate the extent of *Winner’s Curse* bias for each data set.

| GWAS | No. sig. SNPs ($5 \times 10^{-8}$) | Prop. sig. SNPs with *smaller* replication estimate ($5 \times 10^{-8}$) | Prop. sig. SNPs *significantly* overestimated ($5 \times 10^{-8}$) | No. sig. SNPs ($5 \times 10^{-4}$) | Prop. sig. SNPs with *smaller* replication estimate ($5 \times 10^{-4}$) | Prop. sig. SNPs *significantly* overestimated ($5 \times 10^{-4}$) |
| --- | --- | --- | --- | --- | --- | --- |
| **BMI 1** | 6,908 | 0.7135 | 0.2251 | 94,173 | 0.8365 | 0.3386 |
| **BMI 2** | 7,951 | 0.8009 | 0.3202 | 98,351 | 0.8455 | 0.3604 |
| **T2D 1** | 31 | 0.0645<sup>a</sup> | 0.0645 | 5,832 | 0.9830 | 0.8433 |
| **T2D 2** | 76 | 1 | 0.1579 | 5,507 | 0.9951 | 0.8397 |
| **Height 1** | 70,020 | 0.6444 | 0.1829 | 257,000 | 0.6940 | 0.2095 |
| **Height 2** | 70,634 | 0.6824 | 0.1772 | 268,497 | 0.7179 | 0.2406 |

Table 1 details an initial exploration of the six UK Biobank data sets. The number of significant SNPs identified for each data set at two significance thresholds, $5 \times 10^{-8}$ and $5 \times 10^{-4}$ , is provided in the table. The table also contains the proportion of these significant SNPs that were seen to have smaller estimated effect sizes, in terms of absolute value, in their respective replication GWAS. The final column for each threshold provides the proportion of significant SNPs that have significantly overestimated effect sizes. The estimated effect size of a SNP has been defined as significantly overestimated according to Eq (15), but in which the true effect size is replaced by the estimated effect size obtained in the corresponding replication GWAS. These values give an indication of the extent of *Winner's Curse* bias present for each data set and threshold.
<sup>a</sup>When the first T2D data set was used for the discovery GWAS, most of the 31 significant SNPs had larger estimated effect sizes in the replication data set. Discussion of this atypical observation is included in the main text.

#### The problem of Linkage Disequilibrium in real data

Naturally, the results of engagement with real data sets are more complex than those of the simulation study. For example, it was noted that for BMI, in one instance, all significant SNPs which had a *z*-value greater than 15 in the discovery GWAS had association estimates in the replication GWAS which were in fact greater. This observation can be clearly seen in S12 Fig in which *z*-statistics are plotted against estimated bias for each data set, with estimated bias of SNP *i* defined as:

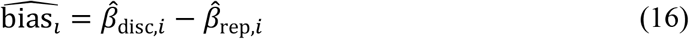

This finding is of course contrary to what is expected. However, these SNPs with *z*-values greater than 15 were all in strong linkage disequilibrium and thus, represented a single independent signal. It can be seen in Table 1 that a similar result was noted when the first T2D data set was used as the discovery GWAS. When using a significance threshold of 5 × 10^-8^, most of the 31 significant SNPs had larger estimated effect sizes in the replication GWAS than in the discovery GWAS. In these cases, we need to be careful not to over-generalize or interpret the results of applying a *Winner’s Curse* correction, given that there may be very few independent association signals at *p* < 5 × 10^-8^.

#### Evaluation of performance at *p* < 5 × 10^-8^

As stated in ‘Materials and methods’, the methods were evaluated using the estimated MSE of SNPs which passed the chosen significance threshold. Using the threshold of 5 × 10^-8^, the estimated MSE for each method and GWAS combination are displayed in Table 2 while Fig 2 provides a corresponding illustration of these values. In this figure, the light blue bar as well as the black dotted horizontal line mark the estimated MSE obtained using the naïve approach, i.e. when no *Winner’s Curse* correction method has been applied and the raw effect estimates are used. This provides a standard to which the performance of each method can be directly compared with, in which it is desired that method application will result in an estimated MSE less than this approach. Similar to the section above describing the results of the simulation study, the poor performance of the conditional likelihood methods is evident.

**Fig 2.**
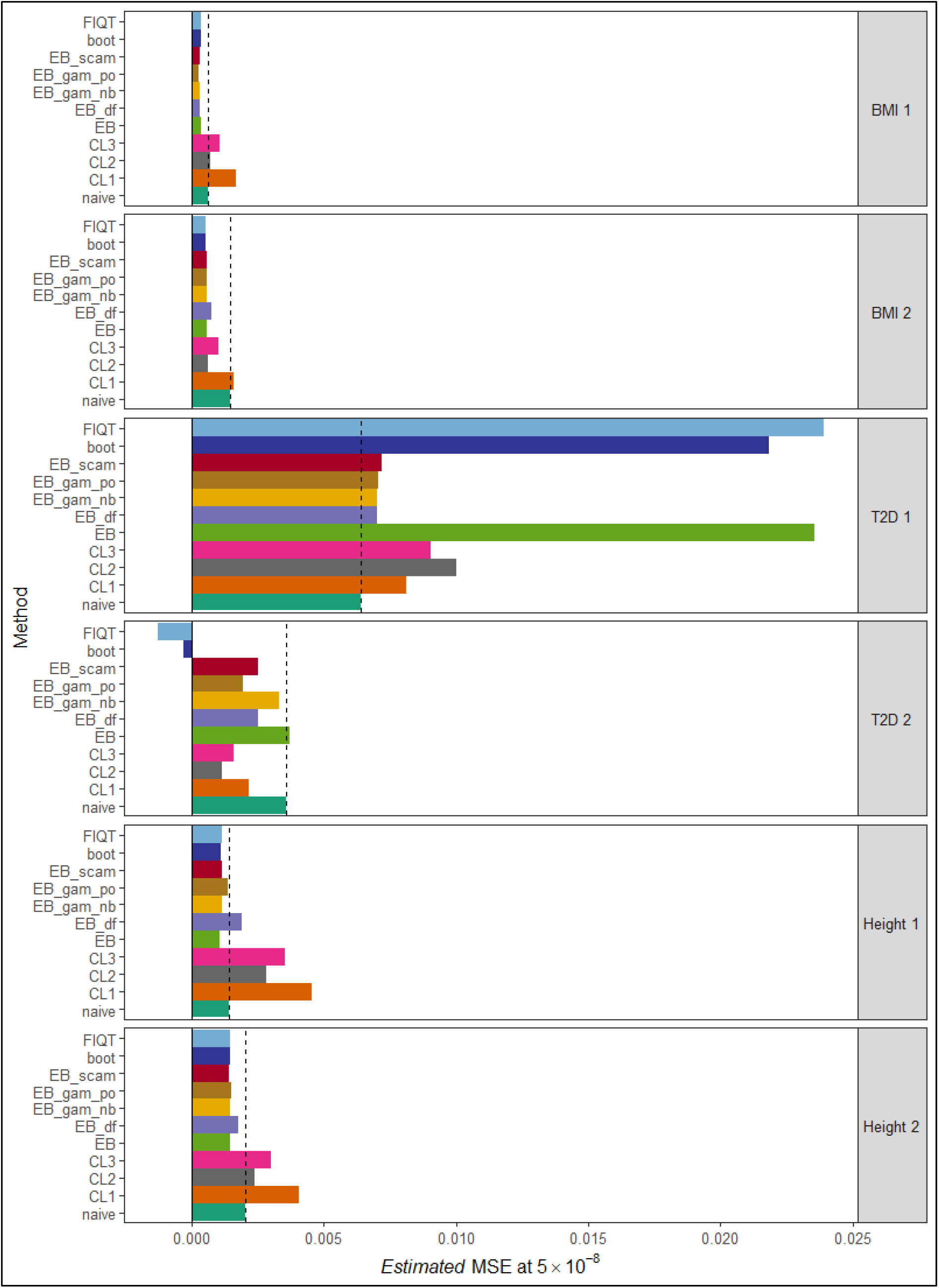
Estimated MSE of significant SNPs at threshold 5 × 10^-8^ for each method and data set. The estimated MSE of significant SNPs at a threshold of 5 × 10^-8^ (x-axis), as defined by Eq (14), is plotted for each correction method (y-axis), for each of the six data sets. The estimated MSE obtained for the naïve approach, when no *Winner’s Curse* correction method is applied, is included, represented by the darker green bar. The dashed black line also represents this value, in order to highlight which methods provide estimated MSE values greater or less than that of the naïve approach. All estimated MSE values plotted here are provided in Table 2.

**Table 2.**
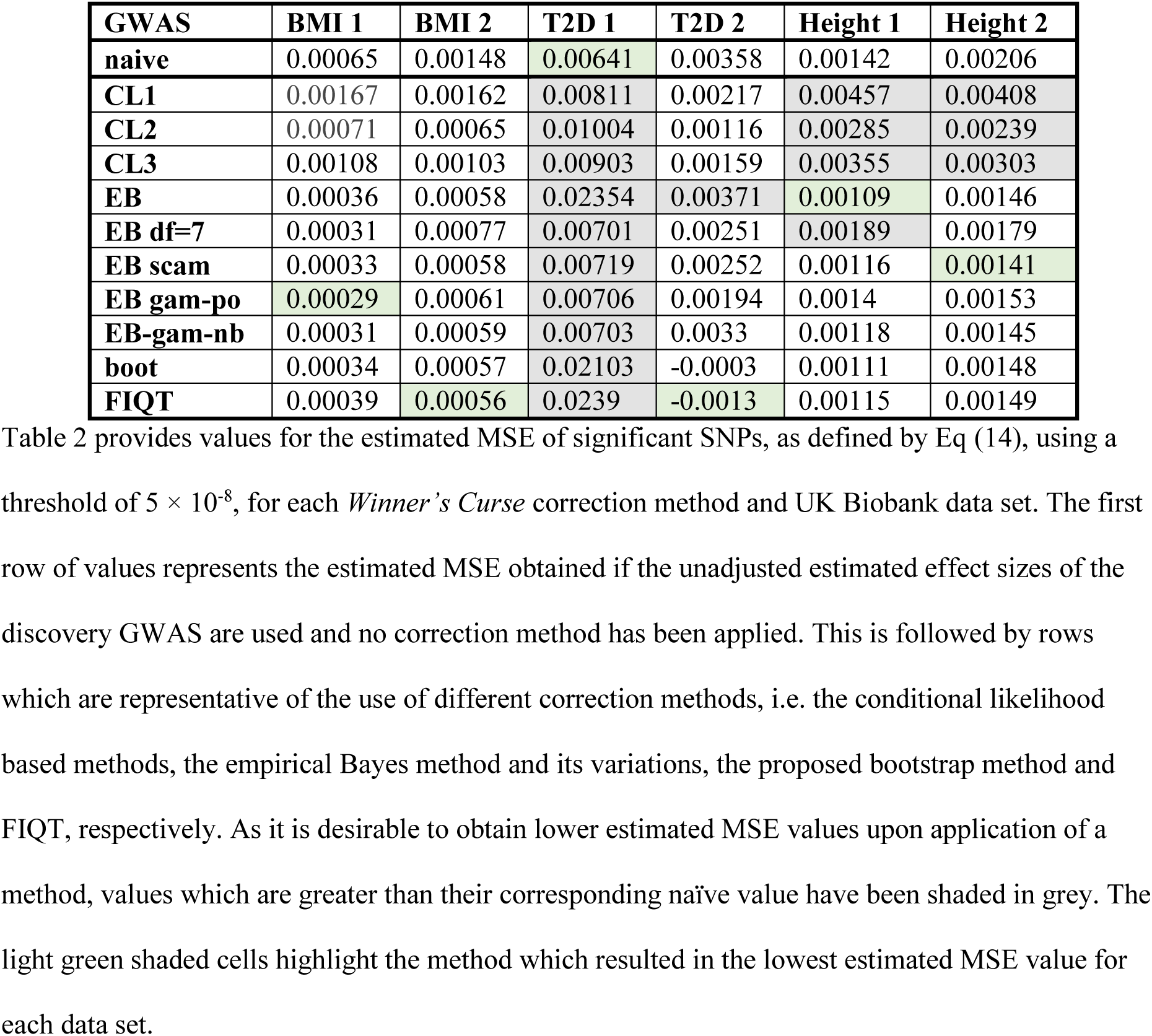
Estimated MSE of significant SNPs at threshold 5 × 10^-8^ for each method and data set.

| GWAS | BMI 1 | BMI 2 | T2D 1 | T2D 2 | Height 1 | Height 2 |
| --- | --- | --- | --- | --- | --- | --- |
| naïve | 0.00065 | 0.00148 | 0.00641 | 0.00358 | 0.00142 | 0.00206 |
| CL1 | 0.00167 | 0.00162 | 0.00811 | 0.00217 | 0.00457 | 0.00408 |
| CL2 | 0.00071 | 0.00065 | 0.01004 | 0.00116 | 0.00285 | 0.00239 |
| CL3 | 0.00108 | 0.00103 | 0.00903 | 0.00159 | 0.00355 | 0.00303 |
| EB | 0.00036 | 0.00058 | 0.02354 | 0.00371 | 0.00109 | 0.00146 |
| EB df=7 | 0.00031 | 0.00077 | 0.00701 | 0.00251 | 0.00189 | 0.00179 |
| EB scam | 0.00033 | 0.00058 | 0.00719 | 0.00252 | 0.00116 | 0.00141 |
| EB gam-po | 0.00029 | 0.00061 | 0.00706 | 0.00194 | 0.0014 | 0.00153 |
| EB-gam-nb | 0.00031 | 0.00059 | 0.00703 | 0.0033 | 0.00118 | 0.00145 |
| boot | 0.00034 | 0.00057 | 0.02103 | -0.0003 | 0.00111 | 0.00148 |
| FIQT | 0.00039 | 0.00056 | 0.0239 | -0.0013 | 0.00115 | 0.00149 |

In 5 out of the 6 independent instances, it was observed that at least one of these methods had a greater estimated MSE than that of the naïve approach.

On real data, the original empirical Bayes method proposed by Ferguson et al. (6) performed poorly, sometimes failing to adjust estimated associations downwards. This observation motivated the proposal of possible modifications as mentioned in ‘Materials and methods’. These suggested improvements, in particular the inclusion of the shape-constrained additive models and the use of the generalized additive model, resulted in slightly more consistent reductions in MSE over significant SNPs. In fact, taking all six data sets into account, it is ‘EB-scam’ and ‘EB-gam-po’, which tend to be the best performing methods, having an average improvement on estimated MSE of greater than 29.4% over the naïve approach.

However, as stated previously, we must be cautious when using the first T2D data set to evaluate methods, and also when using the second T2D data set. The problem with linkage disequilibrium and very few independent signals is common to both data sets. In Fig 2 for the first T2D data set, it is witnessed that all methods result in greater estimated MSE values than the naïve approach, with the original empirical Bayes method, bootstrap and FIQT clearly greatly shrinking the estimated effect sizes of significant SNPs away from those larger replication effect sizes. Therefore, if we exclude these two T2D data sets and re-compute the average improvement in estimated MSE for each method, it is our proposed bootstrap method which is seen to be the dominant method with an average improvement of approximately 40.2%.

#### Evaluation of performance at *p* < 5 × 10^-4^

This evaluation procedure was repeated using a larger significance threshold of 5 × 10^-4^. The results of which can be found summarised in S8 Table and Fig 3. At this threshold, for all 6 data sets, all of the methods produce estimated MSE values less than the naïve approach. Each version of the empirical Bayes method along with the bootstrap and FIQT lead to an average improvement in estimated MSE of between 65 and 70% with the implementation of the empirical Bayes algorithm which incorporates shape constrained additive models (SCAMs) having the greatest average improvement of just over 70%.

**Fig 3.**
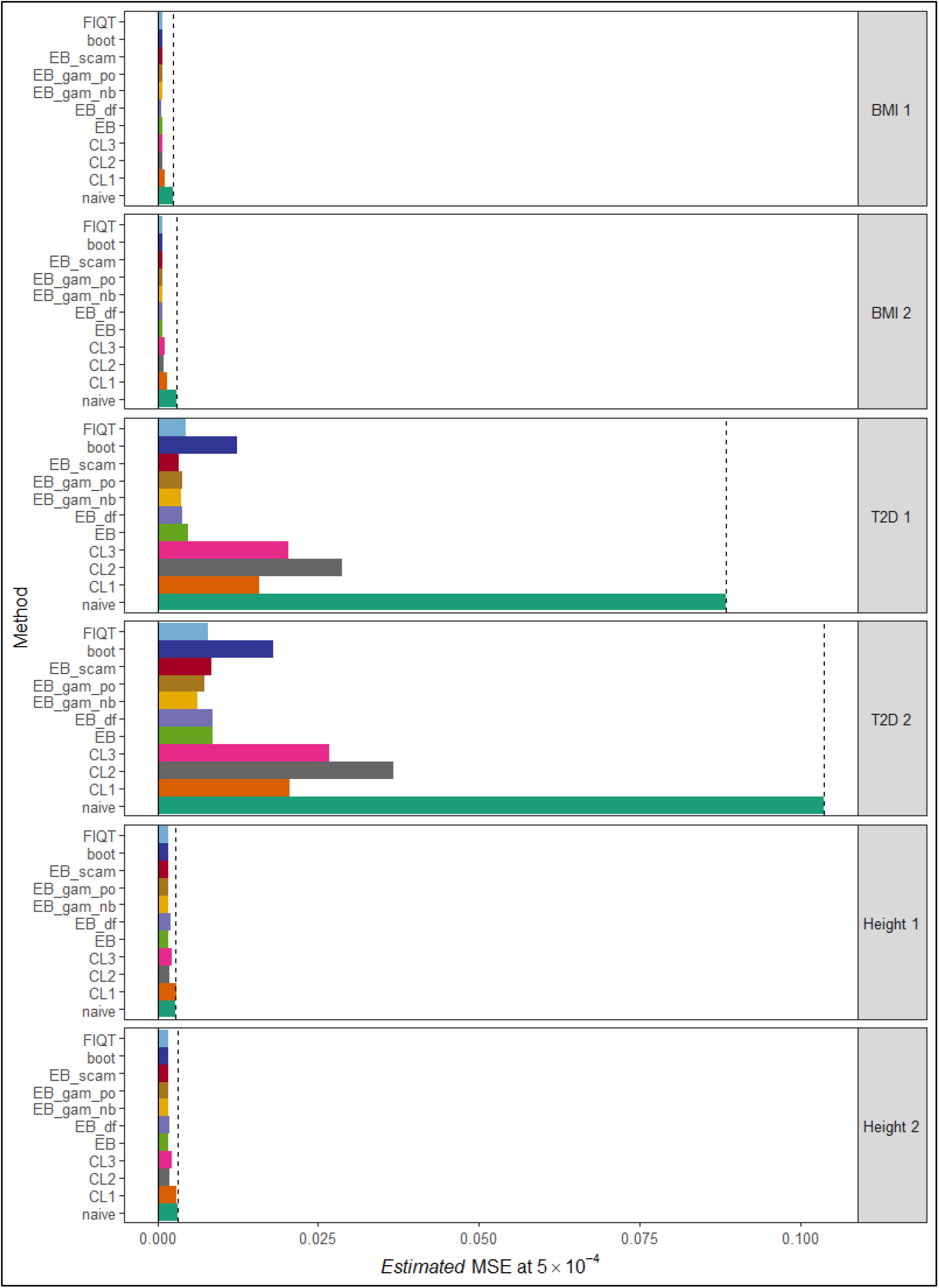
Estimated MSE of significant SNPs at threshold 5 × 10^-4^ for each method and data set. The estimated MSE of significant SNPs at a threshold of 5 × 10^-4^ (x-axis), as defined by Eq (14), is plotted for each correction method (y-axis), for each of the six data sets. The estimated MSE obtained for the naïve approach, when no *Winner’s Curse* correction method is applied, is included, represented by the darker green bar. The dashed black line also represents this value, in order to highlight which methods provide estimated MSE values greater or less than that of the naïve approach. The estimated MSE values plotted here are provided in S6 Table.

## Discussion

In this article, we investigated the problem of *Winner’s Curse* bias which results in the estimated effect sizes of significant SNPs often being greater than their true values. Our work concentrated on methods that could be used to reduce this bias in settings in which only summary statistics of the GWAS that discovered these SNP-trait associations were available. We chose to focus on this particular situation as *Winner’s Curse* correction methods which only require GWAS summary data tend to be very computational efficient and furthermore, this summary data is often much easier to access than individual-level data.

We performed a thorough evaluation and comparison of these methods using both simulated and real data sets. Our simulation study considered a wide range of genetic architectures including data sets in which a simple correlation structure had been imposed on the set of SNPs as well as data sets of independent SNPs. In addition, three UKBB real data sets were used for method evaluation purposes. As well as assessing currently published correction methods, we also explored several possible modifications that could be made in order to improve these methods. In particular, we looked at a number of variations of the empirical Bayes method and proposed an additional approach which uses the parametric bootstrap in order to establish suitable corrections for the estimated effect size of each SNP. The estimated mean squared error (MSE) was chosen as an appropriate metric in order to compare the methods. Due to the notable lack of software for implementation of *Winner’s Curse* correction methods, we developed an R package, ‘winnerscurse’, as an accompaniment to the work described in this paper. This allows users to apply all methods discussed here, as well as the proposed modifications, to their sets of GWAS summary data.

As a first step in both our simulation study and engagement with real data, we computed the proportion of significant SNPs that were significantly overestimated and observed the common trend that as the number of SNPs passing the significance threshold increased, the proportion of those that were significantly overestimated decreased. This aligns with the postulation that as sample sizes increase, *Winner’s Curse* bias becomes less of a concern although it still exists. However, caution must be taken when working with real data sets, especially those of binary traits, in which a very small number of SNPs have been deemed significant at a certain threshold. In this instance, it may be that the significant SNPs are representative of only one or two independent signals. For example, in our first T2D data set, while using a threshold of 5 × 10^-8^, we witnessed 93.5% of significant SNPs having greater replication effect size estimates than those obtained in the discovery GWAS. Fortunately, as sample sizes increase in the future and different diseases have greater numbers of true signals captured by their respective sets of significant SNPs, this issue will only be seen to present itself in rare circumstances.

With respect to method performance, it was clear that the conditional likelihood methods performed poorly as in most instances, especially for *p* < 5 × 10^-8^, these methods resulted in greater values for the estimated MSE among significant SNPs than the naïve approach. The other considered methods behaved much more similarly to the extent that we cannot state that there is a clear advantage of one method over another. Thus, the choice of which method a user should apply to their set of GWAS summary statistics in order to correct for *Winner’s Curse* is dependent on personal preference. However, it is advised that when doing so, the possible limitations of the chosen method are understood well. Notably, the empirical Bayes methods have a clear theoretical advantage, but their performance can be restricted due to inaccurate estimation of the extreme tails of the *z*-statistic distribution. This estimation difficulty is particularly problematic when the existence of strong linkage disequilibrium results in clusters of associations in the tails. These clusters can be falsely detected as local modes in the distribution by automatic fitting algorithms. Some progress on improving estimation in the tails has been made here with the proposal of modifications that employ generalized additive models or shape constrained additive models. However, these adaptations have not resulted in large enough improvements in order to claim objective superiority of the empirical Bayes methods over other approaches such as the bootstrap method or FIQT. In a setting in which the distribution of effect sizes is asymmetric, methods like the empirical Bayes and bootstrap, where the correction rule is not a function of absolute value z-statistics, possess the potential to perform better than FIQT and conditional likelihood methods. In spite of this fact, no tangible evidence of improved performance over FIQT was observed on the real data sets that we examined.

With both our set of simulations and real data analysis, we have aimed to be as comprehensive as possible as it is possible that differing method performance results may occur under differing genetic architectures, but this is an obviously difficult task. Informing these simulations appropriately is particularly challenging, especially when attempting to define the true effect size distribution. However, under the assumption of independent SNPs, we also investigated scenarios which had a bimodal or skewed distribution of effect sizes, as described in the supplementary material. Furthermore, for simulations involving correlated SNPs, we have assumed a very simplistic linkage disequilibrium (LD) structure in which the minor allele frequencies have been simulated independently of this LD structure. In contrast, the use of real data permitted the analysis of method performance in a realistic setting where a large degree of LD exists. However, this was limited to only three UKBB data sets. In the case of the binary trait T2D, it must be noted that due to the very small number of significant SNPs at *p* < 5 × 10^-8^, the results are deemed rather questionable here.

Due to space considerations, *Winner’s Curse* correction methods which require both a discovery and replication GWAS in order to make suitable adjustments to estimated effect sizes have not been examined in this manuscript, even though several of these methods have been implemented in our developed R package, ‘winnerscurse’. Furthermore, computation of standard errors of the adjusted estimated effect sizes have not been considered here.

However, for methods such as the empirical Bayes, bootstrap and FIQT, the R package, ‘winnerscurse’, utilizes the parametric bootstrap in order to obtain these standard errors. This package can also be used to provide confidence intervals for estimated effect sizes which have been corrected for *Winner’s Curse* using the conditional likelihood methods. In two-sample Mendelian randomization, it is known that as this *Winner’s Curse* bias can be present in the estimated SNP-exposure associations, the causal estimate will then suffer from bias. Thus, the *Winner’s Curse* correction methods explored in this paper can also be potentially used as plug-in corrections for two-sample MR. In addition, these methods could prove beneficial in the computation of polygenic risk scores, in order to reduce the effect of *Winner’s Curse* bias there.

## Supporting information

**S1 Fig. Number of significant SNPs at threshold 5 × 10^-8^ plotted against the proportion of those SNPs with significantly overestimated effect sizes, for each simulation setting with a simple correlation structure imposed on the set of SNPs.** For 100 iterations of each of the eight simulation settings, the number of significant SNPs using a significance threshold of 5 × 10^-8^ (x-axis) is plotted against the number of these SNPs with significantly overestimated effect sizes (y-axis). The estimated effect size of a SNP has been defined as significantly overestimated according to Eq (15) in the main text. This figure corresponds to simulation settings in which a simple correlation structure has been imposed on the set of SNPs. These 8 different simulated genetic architectures are defined by combinations of three parameters, sample size *n,* heritability *h^2^* and polygenicity *π.* The parameter values that have been chosen for each simulated scenario are shown in the table, while the legend at the bottom of the plot indicates which colour corresponds to which scenario.

**S2 Fig. *Z*-statistics plotted against bias for each simulation setting, with a simple correlation structure imposed on the set of SNPs.** For a single example of each of the eight simulation settings, *z*-statistics (x-axis) are plotted against bias (y-axis). The *z*-statistic of a SNP is defined as its estimated effect size divided by the standard error of that estimated effect size while the bias of a SNP is defined as its true effect size subtracted from its estimated effect size. This figure corresponds to simulation settings in which a simple correlation structure has been imposed on the set of SNPs. These 8 different simulated genetic architectures are defined by combinations of three parameters, sample size *n*, heritability *h*^2^ and polygenicity *π*. The parameter values that have been chosen for each simulated setting are shown in the subtitle of each plot. In each plot, the dark red dashed vertical line represents the *z*-statistic corresponding to a *p*-value of 5 × 10^-8^ and thus, any points outside these two dark red lines are SNPs with *p*-values passing the genome-wide significance threshold of 5 × 10^-8^. In a similar manner, the light red dashed vertical line represents the greater significance threshold of 5 × 10^-4^. The dark grey points highlight SNPs that have *p*-values less than 5 × 10^-4^ and have significantly overestimated effect sizes, as defined by Eq (15) in the main text.

**S3 Fig. Estimated change in RMSE of significant SNPs at threshold 5 × 10^-4^ for each method and simulation setting, with a simple correlation structure imposed on the set of SNPs.** The estimated change in RMSE of significant SNPs at a threshold of 5 × 10^-4^ (y-axis), as defined by Eq (13) in the main text, is plotted for each correction method (x-axis), for each of the eight simulation settings. This figure corresponds to simulation settings in which a simple correlation structure has been imposed on the set of SNPs.

**S4 Fig. Estimated change in RMSE of significant SNPs at threshold 5 × 10^-8^ for each method and simulation setting, assuming a quantitative trait, independent SNPs and a normal effect size distribution.** The estimated change in RMSE of significant SNPs at a threshold of 5 × 10^-8^ (y-axis), as defined by Eq (13) in the main text, is plotted for each correction method (x-axis), for each of the eight simulation settings with a selection coefficient of zero. This figure corresponds to simulation settings in which it is assumed that the trait of interest is quantitative, SNPs are independent and the effect sizes follow a normal distribution.

**S5 Fig. Estimated change in RMSE of significant SNPs at threshold 5 × 10^-4^ for each method and simulation setting, assuming a quantitative trait, independent SNPs and a normal effect size distribution.** The estimated change in RMSE of significant SNPs at a threshold of 5 × 10^-4^ (y-axis), as defined by Eq (13) in the main text, is plotted for each correction method (x-axis), for each of the eight simulation settings with a selection coefficient of zero. This figure corresponds to simulation settings in which it is assumed that the trait of interest is quantitative, SNPs are independent and the effect sizes follow a normal distribution.

**S6 Fig. Estimated change in RMSE of significant SNPs at threshold 5 × 10^-8^ for each method and simulation setting, assuming a quantitative trait, independent SNPs and a bimodal effect size distribution.** The estimated change in RMSE of significant SNPs at a threshold of 5 × 10^-8^ (y-axis), as defined by Eq (13) in the main text, is plotted for each correction method (x-axis), for each of the eight simulation settings with a selection coefficient of zero. This figure corresponds to simulation settings in which it is assumed that the trait of interest is quantitative, SNPs are independent and the effect sizes follow a bimodal distribution. Note that for this simulation set-up, the alternative variations of the empirical Bayes method have been excluded and only 50 sets of summary statistics were simulated for each setting.

**S7 Fig. Estimated change in RMSE of significant SNPs at threshold 5 × 10^-4^ for each method and simulation setting, assuming a quantitative trait, independent SNPs and a bimodal effect size distribution.** The estimated change in RMSE of significant SNPs at a threshold of 5 × 10^-4^ (y-axis), as defined by Eq (13) in the main text, is plotted for each correction method (x-axis), for each of the eight simulation settings with a selection coefficient of zero. This figure corresponds to simulation settings in which it is assumed that the trait of interest is quantitative, SNPs are independent and the effect sizes follow a bimodal distribution. Note that for this simulation set-up, the alternative variations of the empirical Bayes method have been excluded and only 50 sets of summary statistics were simulated for each setting.

**S8 Fig. Estimated change in RMSE of significant SNPs at threshold 5 × 10^-8^ for each method and simulation setting, assuming a quantitative trait, independent SNPs and a skewed effect size distribution.** The estimated change in RMSE of significant SNPs at a threshold of 5 × 10^-8^ (y-axis), as defined by Eq (13) in the main text, is plotted for each correction method (x-axis), for each of the eight simulation settings with a selection coefficient of zero. This figure corresponds to simulation settings in which it is assumed that the trait of interest is quantitative, SNPs are independent and the effect sizes follow a skewed distribution. Note that for this simulation set-up, the alternative variations of the empirical Bayes method have been excluded and only 50 sets of summary statistics were simulated for each setting.

**S9 Fig. Estimated change in RMSE of significant SNPs at threshold 5 × 10^-4^ for each method and simulation setting, assuming a quantitative trait, independent SNPs and a skewed effect size distribution.** The estimated change in RMSE of significant SNPs at a threshold of 5 × 10^-4^ (y-axis), as defined by Eq (13) in the main text, is plotted for each correction method (x-axis), for each of the eight simulation settings with a selection coefficient of zero. This figure corresponds to simulation settings in which it is assumed that the trait of interest is quantitative, SNPs are independent and the effect sizes follow a skewed distribution. Note that for this simulation set-up, the alternative variations of the empirical Bayes method have been excluded and only 50 sets of summary statistics were simulated for each setting.

**S10 Fig. Estimated change in RMSE of significant SNPs at threshold 5 × 10^-8^ for each method and simulation setting, assuming a binary trait, independent SNPs and a normal effect size distribution.** The estimated change in RMSE of significant SNPs at a threshold of 5 × 10^-8^ (y-axis), as defined by Eq (13) in the main text, is plotted for each correction method (x-axis), for each of the eight simulation settings with a selection coefficient of zero. This figure corresponds to simulation settings in which it is assumed that the trait of interest is binary, SNPs are independent and the effect sizes follow a normal distribution. Note that for this simulation set-up, the alternative variations of the empirical Bayes method have been excluded and only 50 sets of summary statistics were simulated for each setting.

**S11 Fig. Estimated change in RMSE of significant SNPs at threshold 5 × 10^-4^ for each method and simulation setting, assuming a binary trait, independent SNPs and a normal effect size distribution.** The estimated change in RMSE of significant SNPs at a threshold of 5 × 10^-4^ (y-axis), as defined by Eq (13) in the main text, is plotted for each correction method (x-axis), for each of the eight simulation settings with a selection coefficient of zero. This figure corresponds to simulation settings in which it is assumed that the trait of interest is binary, SNPs are independent and the effect sizes follow a normal distribution. Note that for this simulation set-up, the alternative variations of the empirical Bayes method have been excluded and only 50 sets of summary statistics were simulated for each setting.

**S12 Fig. Z-statistics plotted against bias for each real data set.** For each of the six real data sets, *z*- statistics (x-axis) are plotted against estimated bias (y-axis). The *z*-statistic of a SNP is defined as its estimated effect size divided by the standard error of that estimated effect size while the estimated bias of a SNP is defined by Eq (15) in the main text. The title of each plot (A)-(F) indicates which real data set the plot relates to. In each plot, the dark red dashed vertical line represents the *z*-statistic corresponding to a *p*-value of 5 × 10^-8^ and thus, any points outside these two dark red lines are SNPs with *p*-values passing the genome-wide significance threshold of 5 × 10^-8^. In a similar manner, the light red dashed vertical line represents the greater significance threshold of 5 × 10^-4^. The dark grey points highlight SNPs that have *p*-values less than 5 × 10^-4^ and have significantly overestimated effect sizes. The estimated effect size of a SNP has been defined as significantly overestimated according to Eq (15) in the main text, but in which the true effect size is replaced by the estimated effect size obtained in the corresponding replication GWAS.

**S1 Table. The average number and MSE of significant SNPs at two significance thresholds, 5 × 10^-8^ and 5 × 10^-4^, with proportions that indicate the extent of *Winner’s Curse* bias for each simulation scenario.** S1 Table details an initial exploration of the various simulation scenarios. The values provided in the table are averages obtained across 100 simulated sets of summary statistiscs for each scenario. The top portion of the table shows the values of the parameters which define each simulation scenario, i.e. sample size, heritability and polygenicity (proportion of effect SNPs). This table corresponds to simulation settings in which a simple correlation structure has been imposed on the set of SNPs. The number of significant SNPs identified for each scenario at two significance thresholds, 5 × 10^-8^ and 5 × 10^-4^, is provided, as well as the naive MSE of these significant SNPs, as defined by Eq (15) in the main manuscript. The table also contains the proportion of significant SNPs that were seen to have a larger estimated effect size than their true effect size, in terms of absolute value. The final row for each threshold provides the proportion of significant SNPs that have significantly overestimated effect sizes. The estimated effect size of a SNP has been defined as significantly overestimated according to Eq (15) in the main text. This metric gives an indication of the extent of *Winner’s Curse* bias present for each simulation scenario and threshold.

**S2 Table. Estimated change in RMSE of significant SNPs at threshold 5 × 10^-8^ for each method and simulation setting.** S2 Table provides values for the estimated change in RMSE of significant SNPs, as defined by Eq (14) in the main manuscript, for each *Winner’s Curse* correction method and simulation scenario. This table corresponds to simulation settings in which a simple correlation structure has been imposed on the set of SNPs. The top portion of the table shows the values of the parameters which define each simulation scenario, i.e. sample size, heritability and polygenicity (proportion of effect SNPs). As described in the main manuscript, 100 sets of summary statistics were simulated for each scenario and the correction methods were applied to each set. Thus, the values shown in the remaining portion of the table are the average estimated change in RMSE of significant SNPs due to method implementation across each of these 100 sets. As it is the change in RMSE that has been computed, it is desirable to obtain a negative change, i.e. the RMSE computed upon application of the correction method is smaller than that of the naïve approach. Thus, positive values in the table have been shaded in grey, indicating poor performing methods. The light green shaded cells highlight the method which, on average, resulted in the greatest reduction in RMSE for each simulated scenario.

**S3 Table. Estimated change in MSE of significant SNPs at threshold 5 × 10^-8^ for each method and simulation setting.** S3 Table provides values for the estimated change in MSE of significant SNPs, as defined by Eq (14) in the main manuscript, for each *Winner’s Curse* correction method and simulation scenario. This table corresponds to simulation settings in which a simple correlation structure has been imposed on the set of SNPs. The top portion of the table shows the values of the parameters which define each simulation scenario, i.e. sample size, heritability and polygenicity (proportion of effect SNPs). As described in the main manuscript, 100 sets of summary statistics were simulated for each scenario and the correction methods were applied to each set. Thus, the values shown in the remaining portion of the table are the average estimated change in MSE of significant SNPs due to method implementation across each of these 100 sets. As it is the change in MSE that has been computed, it is desirable to obtain a negative change, i.e. the MSE computed upon application of the correction method is smaller than that of the naïve approach. Thus, positive values in the table have been shaded in grey, indicating poor performing methods. The light green shaded cells highlight the method which, on average, resulted in the greatest reduction in MSE for each simulated scenario.

**S4 Table. Estimated relative change in MSE of significant SNPs at threshold 5 × 10^-8^ for each method and simulation setting, with a simple correlation structure imposed on the set of SNPs.** S4 Table provides values for the estimated relative change in MSE of significant SNPs for each *Winner’s Curse* correction method and simulation scenario. This table corresponds to simulation settings in which a simple correlation structure has been imposed on the set of SNPs. The top portion of the table shows the values of the parameters which define each simulation scenario, i.e. sample size, heritability and polygenicity (proportion of effect SNPs). As described in the main manuscript, 100 sets of summary statistics were simulated for each scenario and the correction methods were applied to each set. Thus, the values shown in the remaining portion of the table are the average estimated relative change in MSE of significant SNPs due to method implementation across each of these 100 sets. As it is the relative change in MSE that has been computed, it is desirable to obtain a negative change, i.e. the MSE computed upon application of the correction method is smaller than that of the naïve approach. Thus, positive values in the table have been shaded in grey, indicating poor performing methods. The light green shaded cells highlight the method which, on average, resulted in the greatest relative reduction in MSE for each simulated scenario. As the final column contains the mean of each row, it shows that the bootstrap method has the greatest average estimated relative reduction in MSE. This value of −0.2608 suggests that on average, the bootstrap method improves the MSE of significant SNPs by ≈26.08%.

**S5 Table. Estimated change in RMSE of significant SNPs at threshold 5 × 10^-4^ for each method and simulation setting.** S5 Table provides values for the estimated change in RMSE of significant SNPs, as defined by Eq (14) in the main manuscript, for each *Winner’s Curse* correction method and simulation scenario, when a significance threshold of 5 × 10^-4^ is used. This table corresponds to simulation settings in which a simple correlation structure has been imposed on the set of SNPs. The top portion of the table shows the values of the parameters which define each simulation scenario, i.e. sample size, heritability and polygenicity (proportion of effect SNPs). As described in the main manuscript, 100 sets of summary statistics were simulated for each scenario and the correction methods were applied to each set. Thus, the values shown in the remaining portion of the table are the average estimated change in RMSE of significant SNPs due to method implementation across each of these 100 sets. As it is the change in RMSE that has been computed, it is desirable to obtain a negative change, i.e. the RMSE computed upon application of the correction method is smaller than that of the naïve approach. Thus, positive values in the table have been shaded in grey, indicating poor performing methods. The light green shaded cells highlight the method which, on average, resulted in the greatest reduction in RMSE for each simulated scenario.

**S6 Table. Estimated change in MSE of significant SNPs at threshold 5 × 10^-4^ for each method and simulation setting, with a simple correlation structure imposed on the set of SNPs.** S6 Table provides values for the estimated change in MSE of significant SNPs, as defined by Eq (14) in the main manuscript, for each *Winner’s Curse* correction method and simulation scenario, when a significance threshold of 5 × 10^-4^ is used. This table corresponds to simulation settings in which a simple correlation structure has been imposed on the set of SNPs. The top portion of the table shows the values of the parameters which define each simulation scenario, i.e. sample size, heritability and polygenicity (proportion of effect SNPs). As described in the main manuscript, 100 sets of summary statistics were simulated for each scenario and the correction methods were applied to each set. Thus, the values shown in the remaining portion of the table are the average estimated change in MSE of significant SNPs due to method implementation across each of these 100 sets. As it is the change in MSE that has been computed, it is desirable to obtain a negative change, i.e. the MSE computed upon application of the correction method is smaller than that of the naïve approach. Thus, positive values in the table have been shaded in grey, indicating poor performing methods. The light green shaded cells highlight the method which, on average, resulted in the greatest reduction in MSE for each simulated scenario.

**S7 Table. Estimated relative change in MSE of significant SNPs at threshold 5 × 10^-4^ for each method and simulation setting, with a simple correlation structure imposed on the set of SNPs.**

S7 Table provides values for the estimated relative change in MSE of significant SNPs for each *Winner’s Curse* correction method and simulation scenario, when a significance threshold of 5 × 10^-4^ is used. This table corresponds to simulation settings in which a simple correlation structure has been imposed on the set of SNPs. The top portion of the table shows the values of the parameters which define each simulation scenario, i.e. sample size, heritability and polygenicity (proportion of effect SNPs). As described in the main manuscript, 100 sets of summary statistics were simulated for each scenario and the correction methods were applied to each set. Thus, the values shown in the remaining portion of the table are the average estimated relative change in MSE of significant SNPs due to method implementation across each of these 100 sets. As it is the relative change in MSE that has been computed, it is desirable to obtain a negative change, i.e. the MSE computed upon application of the correction method is smaller than that of the naïve approach. Thus, positive values in the table have been shaded in grey, indicating poor performing methods. The light green shaded cells highlight the method which, on average, resulted in the greatest relative reduction in MSE for each simulated scenario. As the final column contains the mean of each row, it shows that the original empirical Bayes method has the greatest average estimated relative reduction in MSE, when a significance threshold of 5 × 10^-4^ is used. This value of −0.3338 suggests that on average, this form of the empirical Bayes method improves the MSE of significant SNPs by ≈33.38%.

**S8 Table. Estimated MSE of significant SNPs at threshold 5 × 10^-4^ for each method and data set.**

S8 Table provides values for the estimated MSE of significant SNPs, as defined by Eq (14) in the main manuscript, using a threshold of 5 × 10^-4^, for each *Winner’s Curse* correction method and UK Biobank data set. The first row of values represents the estimated MSE obtained if the unadjusted estimated effect sizes of the discovery GWAS are used and no correction method has been applied. This is followed by rows which are representative of the use of different correction methods, i.e. the conditional likelihood based methods, the empirical Bayes method and its variations, the proposed bootstrap method and FIQT, respectively. As it is desirable to obtain lower estimated MSE values upon application of a method, values which are greater than their corresponding naïve value have been shaded in grey. The light green shaded cells highlight the method which resulted in the lowest estimated MSE value for each data set.

**S1 File. Text Supplement.** This file contains a more detailed description of the various proposed modifications to the empirical Bayes method, the simulation process and the evaluation of method performance using simulated data sets of independent SNPs.

