## Supplementary File 1 for "Review and further developments in statistical corrections for Winner’s Curse in genetic association studies"

***** Corresponding author

### Contents

**Modifications to the Empirical Bayes method** 1

**Simulation details** 2

- **Quantitative trait with a correlation structure imposed** 2
- **Quantitative trait with independence assumed** 4
- **Binary trait with independence assumed** 5

**Evaluation of performance assuming independent SNPs** 8

### Modifications to the Empirical Bayes method

As mentioned in the main manuscript, our work incorporated the exploration of several variations of the empirical Bayes method. We first adapted the method by simply altering the minimum-BIC estimated basis function of the natural cubic spline so that the boundary knots were no longer designated to be the most extreme *z*-statistics. Instead, the lower boundary knot is defined as the $10^{th}$ *z*-statistic when the *z*-statistics lie in increasing order while the upper boundary knot is the $10^{th}$ *z*-statistic when the *z*-statistics have been arranged in decreasing order. With *z*-statistics labelled in increasing order such that $z_1 < z_2 < \cdots < z_N$ , this constraint ensures that the estimated log *p(z)* is linear beyond these boundary knots, i.e. below $z_{10}$ and above $z_{N\text{-}10}$.

Following this, together with the above modification, the exclusion of the utilization of BIC for model selection purposes was considered which limited the number of knots in the spline. We reverted back to abiding by Efron's original specification of setting the degrees of freedom to 7 (11). The motivation for this choice stemmed from evaluating the performance of the empirical Bayes method with real data sets in which it was seen that the BIC approach generally selected a large number of basis functions, resulting in severe overfitting. The reason for this being perhaps due to these model selection criteria not accounting for the presence of strong linkage disequilibrium in real data sets.

This observation was also responsible for the investigation of models that enforced additional constraints on the shape of the estimated log density function. We assessed a variation of the empirical Bayes method in which the gam function in the R package mgcv (12) was employed. The gam function fits a generalized additive model (GAM), with smoothness estimation integrated in the fitting process. The smoothing parameters are selected by means of generalized cross-validation (GCV). Two uses of this function were investigated, one in which the distribution family was specified as poisson and the other in which the identification of the family as negative binomial took place. The negative binomial is considered to be the more realistic choice here as it accounts for the overdispersion that is typically found in this form of count data.

In addition, the scam function in the R package scam (13) was used to apply two shape constrained additive models (SCAMs) to the dataset at hand. This action imposes the restriction that only monotone increasing smooths can be attained for *z*-statistics greater than or equal to zero. Alternatively, for negative *z*-statistics, it results in smooths which are monotone decreasing.

### Simulation details

#### Quantitative trait with a correlation structure imposed

The following steps were executed in order to obtain an estimated effect size, $\hat{\beta}_i$ and corresponding standard error, se($\hat{\beta}_i$) for SNP $i$, $i = 1, \ldots, N$, for a quantitative trait in which a simple correlation structure has been imposed on the set of SNPs and the true effect sizes follow a normal distribution. $N$ is the total number of SNPs, which has been fixed at $N$ = 1,000,000 for this simulation study.

1) A value is provided for $\pi$, the proportion of the total SNPs which are truly associated with the trait in question. This determines the number of effect SNPs $K = \pi \cdot N$, $1 \leq K \leq N$, forming a polygenic background.
2) A minor allele frequency is attained for SNP $i$ using the uniform distribution $\text{maf}_i \sim U[0.01, 0.5]$.
3) Let us assume that the true effect sizes are $\boldsymbol{b} = b_1, \ldots, b_N$. We then let $\boldsymbol{X}$ be an $n \times N$ matrix of genotypes, assumed to be mean centred, that is for each column $i$: $\sum_j X_{j,i} = 0$. Recall that $n$ represents the sample size or total number of individuals. We assume that genotypes $\boldsymbol{X}$ affects the $n \times 1$ response vector $\boldsymbol{Y}$ through the following linear model:

$$\boldsymbol{Y} = \boldsymbol{X}\boldsymbol{b} + \varepsilon \qquad \text{(S1)}$$

where $\varepsilon$ is a vector of independent zero mean normally distributed errors, i.e. it is assumed that $\text{var}(\varepsilon) = \sigma^2\boldsymbol{I}$ where $\boldsymbol{I}$ denotes the $n \times n$ identity matrix. Now, let $\boldsymbol{D} = \text{Diag}(d_1, \ldots, d_N)$ where $d_i = \sum_j X^2_{j,i}$. Using $E(X^2_i) = \sum_j X^2_{j,i} / n$, an approximation via the Law of Large Numbers, and the identity $\text{var}(X_i) = E(X^2_i) - (E(X_i))^2$, we have:

$$d_i = n \cdot \text{var}(X_i) = n \cdot 2 \cdot \text{maf}_i(1 - \text{maf}_i) \qquad \text{(S2)}$$

as each column of genotypes is assumed to be mean centred, $E(X_i) = 0$ for each $i$ and the variance of each SNP $i$, $\text{var}(X_i)$, is assumed to take the form $\text{var}(X_i) = 2 \cdot \text{maf}_i(1-\text{maf}_i)$.

With the above definition for $\boldsymbol{D}$, the regression coefficients, $\hat{\beta}_1, \ldots, \hat{\beta}_N$ from the marginal regression of $Y$ on each SNP $X_i$ can be computed via the vector equation:

$$\hat{\beta} = \left(\boldsymbol{X^T X}\right)^{-1}\boldsymbol{X^T Y} = \boldsymbol{D}^{-1}\boldsymbol{X^T Y}\,. \qquad \text{(S3)}$$

Conditionally on the genotype matrix $\boldsymbol{X}$, these estimated regression coefficients have variance-covariance matrix:

$$\textbf{cov}\left(\hat{\beta}\right) = \boldsymbol{D}^{-1}\boldsymbol{X^T X D}^{-1}\sigma^2 = \boldsymbol{D}^{-\frac{1}{2}}\boldsymbol{D}^{-\frac{1}{2}}\boldsymbol{X^T X D}^{-\frac{1}{2}}\boldsymbol{D}^{-\frac{1}{2}}\sigma^2 \approx \boldsymbol{D}^{-\frac{1}{2}}\boldsymbol{R D}^{-\frac{1}{2}}\sigma^2 \qquad \text{(S4)}$$

where $\boldsymbol{R}$ is the $N \times N$ LD matrix of inter-genotype correlations, which should approximately equal the empirical correlation matrix $\boldsymbol{D}^{-½}\boldsymbol{X^T X}\,\boldsymbol{D}^{-½}\,\boldsymbol{D}^{-½}$. Letting $\boldsymbol{SE}$ be the $N \times N$ diagonal matrix with element $i$ equal to $\text{se}\left(\hat{\beta}_i\right) = \sigma / \sqrt{\sum_{j=1}^{n} X^2_{j,i}}$, the final expression could instead be written as: $\textbf{cov}\left(\hat{\beta}\right) = (\boldsymbol{SE})^{-\frac{1}{2}}\boldsymbol{R}(\boldsymbol{SE})^{-\frac{1}{2}}$.

Finally, these estimated associations are not necessarily unbiased for the true causal effects $b_1, \ldots, b_N$. Instead they are unbiased for:

$$E(\hat{\beta}) = \boldsymbol{D}^{-1}\boldsymbol{X}^{\boldsymbol{T}}E(\boldsymbol{Y}) = \boldsymbol{D}^{-1}\boldsymbol{X}^{\boldsymbol{T}}E(\boldsymbol{Xb}+\varepsilon) = \boldsymbol{D}^{-1}\boldsymbol{X}^{\boldsymbol{T}}\boldsymbol{Xb}$$
$$= \boldsymbol{D}^{-\frac{1}{2}}\boldsymbol{D}^{-\frac{1}{2}}\boldsymbol{X}^{\boldsymbol{T}}\boldsymbol{X}\boldsymbol{D}^{-\frac{1}{2}}\boldsymbol{D}^{\frac{1}{2}}\boldsymbol{b} \approx \boldsymbol{D}^{-\frac{1}{2}}\boldsymbol{R}\boldsymbol{D}^{\frac{1}{2}}\boldsymbol{b} \quad \text{(S5)}$$

Multivariate normality of $\hat{\beta}$ is inherited from the assumption that $\varepsilon$ is normally distributed. In summary, conditional on the centred genotype matrix $\boldsymbol{X}$:

$$\hat{\beta} \sim N\left(\boldsymbol{D}^{-\frac{1}{2}}\boldsymbol{R}\boldsymbol{D}^{\frac{1}{2}}\boldsymbol{b}, \boldsymbol{D}^{-\frac{1}{2}}\boldsymbol{R}\boldsymbol{D}^{-\frac{1}{2}}\sigma^2\right) \quad \text{(S6)}$$

4) Assuming that the true effect sizes, $b_i$, $i = 1, \ldots, K$, of associated SNPs follow a Gaussian distribution with mean 0, $b_i$ is then sampled for SNP $i$, $i = 1, \ldots, K$, from the distribution $b_i \sim N(0, [2\text{maf}_i(1-\text{maf}_i)]^S)$. Recall that $S$ is the value of the selection coefficient, which is fixed at 0 for these simulations. Non-effect SNP $i$, $i = K+1, \ldots, N$, is simply assigned the null true effect size, $b_i = 0$.
5) Defining the heritability $h^2$ as the proportion of phenotypic variation, var($Y$), that is explained by all SNPs, var($Y$) can be computed as

$$\text{var}(Y) = \frac{\sum_{i=1}^{K} 2\text{maf}_i(1-\text{maf}_i)\cdot b_i^2}{h^2} \quad \text{(S7)}$$

and following this, the true effect sizes are scaled giving $b_i = \frac{b_i}{\sqrt{\text{var}(Y)}}$ for $i = 1, \ldots, N$, in order to ensure a phenotype with variance 1, i.e. $\sigma^2 = 1$.
6) Next, using the R function sample, $K$ random positions between 1 and $N$ are chosen for the effect SNPs and the vectors containing the values of true effect size, $b_i$ and minor allele frequency, $\text{maf}_i$ are adjusted accordingly.
7) In order to reduce computation time, it is assumed that the same linkage disequilibrium structure exists in independent blocks of 100 SNPs. Therefore, for each block of 100 SNPs, the estimated effect sizes, $\hat{\beta}_i$ are simulated using Eq (S6). As we have already scaled $b_i$ to ensure the phenotype has variance 1, we have $\sigma^2 = 1$. The matrix $\boldsymbol{D}$ is a diagonal 100 × 100 matrix. Thus, using Eq (S2), $\boldsymbol{D}^{-1/2}$ is a similar diagonal matrix with $d_i^{-1/2} = \frac{1}{\sqrt{n\cdot 2\cdot \text{maf}_i(1-\text{maf}_i)}}$ and $\boldsymbol{D}^{1/2}$ has diagonal entries $d_i^{1/2} = \sqrt{n\cdot 2\cdot \text{maf}_i(1-\text{maf}_i)}$. The challenge here is to choose a suitable matrix for $\boldsymbol{R}$, the 100 × 100 LD matrix of inter-genotype correlations. For simplicity, we have chosen $\boldsymbol{R}$ to be of the following format:

$$\boldsymbol{R} = \begin{pmatrix} 1 & \rho & \rho^2 & \rho^3 & \cdots & \rho^{99} \\ \rho & 1 & \rho & \rho^2 & \cdots & \rho^{98} \\ \rho^2 & \rho & 1 & \rho & \cdots & \rho^{97} \\ \vdots & \vdots & \vdots & \vdots & \ddots & \vdots \\ \rho^{99} & \rho^{98} & \rho^{97} & \rho^{96} & \cdots & 1 \end{pmatrix} \quad \text{(S8)}$$

The task is now to choose a suitable value for $\rho$. In Bosch et al. (16), it can be seen in Figure 1a that the average $r^2$ at a distance of 5,000 bases is just over 0.4 in Europeans. Using our UKBB data set, we computed the median distance between SNPs and obtained a value of 189 bases. This would suggest that there exist an average of approximately 26 SNPs per 5,000 bases and thus, a rough estimate of an appropriate value for $\rho$ may be computed as follows:

$$\rho^{26} \approx \sqrt{0.4} \approx 0.63245 \rightarrow \rho \approx 0.9825. \quad \text{(S9)}$$

Within each block of 100 SNPs, we then simulated values for $E(\hat{\beta})$, as defined by Eq (S5), and $\hat{\beta}$. We obtained $\hat{\beta}$ using the R function rnorm with $\hat{\beta} = E(\hat{\beta}) + \boldsymbol{R}^{\frac{1}{2}}\boldsymbol{D}^{-\frac{1}{2}} \cdot$ rnorm(100). The standard errors for each SNP, $\text{se}(\hat{\beta}_i)$ were easily obtained from the diagonal entries of $\boldsymbol{D}^{-½}$.

8) The above process thus provided values for $E(\hat{\beta}_i), \hat{\beta}_i$ and $\text{se}(\hat{\beta}_i)$ for each SNP $i$ = 1, ..., $N$, in which the same LD structure described by the simple matrix $\boldsymbol{R}$ has been imposed for each independent block of 100 SNPs.

#### Quantitative trait with independence assumed

Similar steps were followed in order to obtain an estimated effect size, $\hat{\beta}_i$ and corresponding standard error, $\text{se}(\hat{\beta}_i)$ for SNP $i$, $i$ = 1, ..., $N$, in which a quantitative trait was considered, it was assumed that SNPs were independent and effect sizes follow a normal distribution.

1) A polygenic background of $K = \pi \cdot N$, $1 \leq K \leq N$, effect SNPs is formed, in which $\pi$ is the proportion of the total SNPs which are truly associated with the trait in question.
2) A minor allele frequency is attained for SNP $i$ using the uniform distribution $\text{maf}_i \sim U[0.01, 0.5]$.
3) Assuming that the true effect sizes, $\beta_i$ of associated SNPs follow a Gaussian distribution with mean 0, $\beta_i$ is sampled for SNP $i$, $i$ = 1, ..., $K$, from the distribution $\beta_i$ ~ ~ $N(0, [2\text{maf}_i(1- \text{maf}_i)]^S)$. Non-effect SNP $i$, $i = K$+1, ..., $N$, is simply assigned the null true effect size, $\beta_i = 0$.
   Defining the heritability $h^2$ as the proportion of phenotypic variation, var($Y$), that is explained by all SNPs, var($Y$) is computed in the same manner as Eq (S7) but with $b_i$ replaced by $\beta_i$. Following this, the true effect sizes are scaled by dividing each by the square root of var($Y$) in order to ensure a phenotype with variance 1.
4) For a single SNP $i$, it is assumed that the underlying relationship between $y_j$, a numerical measurement of the trait of individual $j$ and $x_j \in \{0,1,2\}$, the number of minor alleles that individual $j$ has at SNP $i$, is described by the simple linear model

   $$y_j = \beta_0 + \beta_1 x_j + \varepsilon_j \quad \text{(S10)}$$

   for $j$ = 1, ...., $n$. In this equation, $\beta_1$ is recognised as the effect size of SNP $i$, i.e. $\beta_1 = \beta_i$. Using the properties of a linear model, we obtain

   $$\text{se}(\hat{\beta}_i) = \sqrt{\frac{1 - 2\text{maf}_i(1 - \text{maf}_i) \cdot \beta_i^2}{(n-2) \cdot 2\text{maf}_i(1 - \text{maf}_i)}} \quad \text{(S11)}$$

   for each SNP $i$, $i$ = 1, ..., $N$.

5) Finally, assuming that the effect size of each SNP follows a Gaussian distribution with mean $\beta_i$ and standard deviation $\text{se}(\hat{\beta}_i)$, an estimated effect size, $\hat{\beta}_i$ is simulated for each SNP $i$, $i$ = 1, ..., $N$, i.e. $\hat{\beta}_i \sim N\left(\beta_i,\ \text{se}(\hat{\beta}_i)\right)$.

For a bimodal distribution of effect sizes, summary statistics are simulated in the exact same manner but with step 3) above replaced by:

1) It is assumed that the true effect sizes, $\beta_i$ of half of the associated SNPs follow a Gaussian distribution with mean 2.5 while the true effect sizes, $\beta_i$ of the other half follow a Gaussian distribution with mean 0. Thus, $\beta_i$ is sampled for SNP $i$, $i$ = 1, ..., $K/2$, from the distribution $\beta_i \sim \sim N(2.5, [2\text{maf}_i(1-\text{maf}_i)]^S)$ and for SNP SNP $i$, $i = K/2 + 1, ..., K$, $\beta_i$ is sampled from the distribution $\beta_i \sim \sim N(0, [2\text{maf}_i(1-\text{maf}_i)]^S)$. As above, non-effect SNP $i$, $i = K+1, ..., N$, is assigned the null true effect size, $\beta_i = 0$.

Similarly, for a skewed distribution of effect sizes, step 3) is altered and takes the form of:

3) It is assumed that the true effect sizes, $\beta_i$ of 10% of the associated SNPs follow a negative exponential distribution with rate $([2\text{maf}_i(1-\text{maf}_i)]^S)^{-1/2}$ $\frac{1}{\sqrt{[2\text{maf}_i(1-\text{maf}_i)]^S}}$ while the true effect sizes, $\beta_i$ of the other 90% follow an exponential distribution with the same rate. Thus, $\beta_i$ is sampled for SNP $i$, $i$ = 1, ..., $K/10$, from the distribution $\beta_i \sim -\text{Exp}(\frac{1}{\sqrt{[2\text{maf}_i(1-\text{maf}_i)]^S}})$ and for SNP SNP $i$, $i = K/10 + 1, ..., K$, $\beta_i$ is sampled from the distribution $\beta_i \sim \text{Exp}(\frac{1}{\sqrt{[2\text{maf}_i(1-\text{maf}_i)]^S}})$. As above, non-effect SNP $i$, $i = K+1$, ..., $N$, is assigned the null true effect size, $\beta_i = 0$.

#### Binary trait with independence assumed

Maintaining a normal effect size distribution and a set of independent SNPs, the following steps were executed in order to obtain an estimated effect size, $\hat{\beta}_i$ and corresponding standard error, $\text{se}(\hat{\beta}_i)$ for SNP $i$, $i$ = 1, ..., $N$, for a binary trait with disease prevalence of 0.1.

1) A polygenic background of $K = \pi \cdot N$, $1 \leq K \leq N$, effect SNPs is formed, in which $\pi$ is the proportion of the total SNPs which are truly associated with the trait in question.
2) A minor allele frequency is attained for SNP $i$ using the uniform distribution $\text{maf}_i \sim U[0.01, 0.5]$.
3) Assuming that the true effect sizes, $\beta_i$ of associated SNPs follow a Gaussian distribution with mean 0, $\beta_i$ is sampled for SNP $i$, $i$ = 1, ..., $K$, from the distribution $\beta_i \sim \sim N(0, [2\text{maf}_i(1-\text{maf}_i)]^S)$. Non-effect SNP $i$, $i = K+1, ..., N$, is simply assigned the null true effect size, $\beta_i = 0$.
4) Defining the heritability $h^2$ of a binary trait as:

$$h^2 = \frac{\sum_{i=1}^{k} 2\text{maf}_i(1-\text{maf}_i) \cdot \beta_i^2}{1.6^2 + \sum_{i=1}^{k} 2\text{maf}_i(1-\text{maf}_i) \cdot \beta_i^2} \tag{S12}$$

the true effect sizes are re-scaled giving

$$\beta_i = \beta_i \cdot \sqrt{\frac{1.6^2 h^2}{(1-h^2)\sum_{i=1}^{k} 2\text{maf}_i(1-\text{maf}_i) \cdot \beta_i^2}} \quad \text{(S13)}$$

for $i = 1, ..., N$.

5) For a single SNP $i$, it is assumed that the underlying relationship between $y_j$, a numerical measurement of the trait of individual $j$ and $x_j \in \{0,1,2\}$, the number of minor alleles that individual $j$ has at SNP $i$, is described by the logistic model:

$$\text{logit}(\text{P}(y_j = 1|x_j)) = \beta_0 + \beta_1 x_j + \varepsilon_j \quad \text{(S14)}$$

for $j = 1, ..., n$. In this equation, $\beta_1$ is recognised as the effect size of SNP $i$, i.e. $\beta_1 = \beta_i$. For SNP $i$, as we have simulated a value for both its minor allele frequency $\text{maf}_i$ and true effect size $\beta_i = \beta_1$, we can obtain a corresponding value for $\beta_0$ using the fact that we have chosen the disease prevalence to be 0.1, i.e. $P(Y = 1) = 0.1$. Therefore, as $P(Y = 1) = \sum P(Y = 1 \mid X)\, P(X)$ and $P(Y = 1|X) = \frac{e^{\beta_0+\beta_1 X}}{1+e^{\beta_0+\beta_1 X}}$, for SNP $i$, we have:

$$\text{maf}_i^2 \cdot \frac{e^{\beta_0+2\beta_1}}{1+e^{\beta_0+2\beta_1}} + 2\text{maf}_i(1-\text{maf}_i) \cdot \frac{e^{\beta_0+\beta_1}}{1+e^{\beta_0+\beta_1}} + (1-\text{maf}_i)^2 \cdot \frac{e^{\beta_0}}{1+e^{\beta_0}} = 0.1 \quad \text{(S15)}$$

Solving this equation then provides a value for $\beta_0$, given values for $\text{maf}_i$ and $\beta_1$. Now, the aim is to obtain a value for $\text{se}(\hat{\beta}_i) = \text{se}(\hat{\beta}_1)$ for each SNP $i$. Firstly, let us denote the maximum likelihood estimated logistic regression coefficient vector, from regressing the $n \times 1$ response vector $\boldsymbol{Y}$ on the $n \times 1$ genotype vector $\boldsymbol{X}$, as $\widehat{\boldsymbol{\beta}} = \begin{pmatrix} \hat{\beta}_0 \\ \hat{\beta}_1 \end{pmatrix}$, which can be represented as:

$$\widehat{\boldsymbol{\beta}} = \left(\boldsymbol{X}^T \widehat{\boldsymbol{W}} \boldsymbol{X}\right)^{-1} \boldsymbol{X}(\boldsymbol{Y} - \widehat{\boldsymbol{p}}) \quad \text{(S16)}$$

Here,

$$\widehat{\boldsymbol{W}} = \begin{pmatrix} \hat{p}_1(1-\hat{p}_1) & 0 & \cdots & 0 \\ 0 & \hat{p}_2(1-\hat{p}_2) & \cdots & 0 \\ \vdots & \vdots & \ddots & \vdots \\ 0 & 0 & \cdots & \hat{p}_n(1-\hat{p}_n) \end{pmatrix} \quad \text{(S17)}$$

$$\hat{p}_j = \frac{e^{\hat{\beta}_0+\hat{\beta}_1 x_j}}{1+e^{\hat{\beta}_0+\hat{\beta}_1 x_j}} \quad \text{(S18)}$$

and

$$\widehat{\boldsymbol{p}} = (\hat{p}_1, \dots, \hat{p}_n)^T \quad \text{(S19)}$$

It is well known that $\text{var}(\widehat{\boldsymbol{\beta}}) \sim (\boldsymbol{X}^T \widehat{\boldsymbol{W}} \boldsymbol{X})^{-1}$. This can be re-expressed as:

$$X^T\widehat{W}X = \begin{pmatrix} \sum_{j=1}^{n} \hat{p}_j(1-\hat{p}_j) & \sum_{j=1}^{n} X_j\hat{p}_j(1-\hat{p}_j) \\ \sum_{j=1}^{n} X_j\hat{p}_j(1-\hat{p}_j) & \sum_{j=1}^{n} X_j^2\hat{p}_j(1-\hat{p}_j) \end{pmatrix} \sim (n)\begin{pmatrix} \mathrm{E}\big(P(1-P)\big) & \mathrm{E}\big(GP(1-P)\big) \\ \mathrm{E}\big(GP(1-P)\big) & \mathrm{E}\big(G^2P(1-P)\big) \end{pmatrix} \quad \text{(S20)}$$

where $P$ is the random variable that takes value $\mathrm{P}(Y = 1|X) = \frac{e^{\beta_0+\beta_1 X}}{1+ e^{\beta_0+\beta_1 X}}$ for genotype $X$, and $X$ is the binomial distribution for genotype assuming Hardy Weinberg Equilibrium. Therefore, using the above, we obtain the inverse of $X^T\widehat{W}X$. Taking the square root of the value in the second row and second column of this matrix provides $\mathrm{se}(\hat{\beta}_1)$. This process is repeated for each SNP $i$, $i = 1, ..., N$.

6) Finally, assuming that the effect size of each SNP follows a Gaussian distribution with mean $\beta_i$ and standard deviation $\mathrm{se}(\hat{\beta}_i)$, an estimated effect size, $\hat{\beta}_i$ is simulated for each SNP $i$, $i = 1, ..., N$, i.e. $\hat{\beta}_i \sim N\left(\beta_i,\ \mathrm{se}(\hat{\beta}_i)\right)$.

#### Evaluation of performance assuming independent SNPs

Surprisingly, in this instance in which SNPs are assumed to be independent, with respect to the evaluation metric 'change in RMSE of significant SNPs' the simulations suggest that many of the investigated *Winner's Curse* methods tend to break down, or no longer make improvements when the proportion of effect SNPs is 0.001. In these simulations, a replication approach, which selects significant SNPs using a discovery GWAS and then employs a replication GWAS of the same size to obtain unbiased association estimates for these SNPs, was also considered. This can be viewed as acting as a form of benchmark for the other methods. Both the empirical Bayes method and 'EB-gam-nb' perform very similar to this replication approach, as can be seen in S4 Fig. Under the assumption that SNPs are independent, this observation supports the use of these two methods to adjust for *Winner's Curse* bias, particularly when a replication GWAS is not available. The consistency of the methods is what makes them stand out. Unlike the bootstrap, FIQT and other variations of the empirical Bayes method, applying 'EB' or 'EB-gam-nb' rarely results in an increase in the RMSE over all significant SNPs.

It is surprising to see the empirical Bayes method which uses two shape constrained additive models (SCAMs) perform extremely poorly when the sample size is 300,000 and the proportion of effect SNPs 0.001. This method was seen to perform well with real data and when a correlation structure was imposed on simulated data. When the proportion of effect SNPs is 0.01, the proposed bootstrap method for summary statistics performs in a comparable manner to both empirical Bayes and the replication method. However, the bootstrap method ceases to perform as well at obtaining less biased association estimates when the proportion of effect SNPs is reduced to 0.001. That said, it is still seen to perform competitively with respect to other currently published methods. Just as was observed in the simulations with linkage disequilibrium, described in the main manuscript, the conditional likelihood methods tend to perform poorly compared to the other methods. The average percentage improvement in estimated MSE across all scenarios with a selection coefficient of zero was 11.8% for 'EB-gam-nb' and 23.8% for 'EB', although this average metric was negative for some methods, such as the conditional likelihood based approaches, indicating increased inaccuracy from applying *Winner's Curse* corrections.

Imposing a threshold of $5 \times 10^{-4}$, it is still the original empirical Bayes method which seems to be the best, as can be seen in S5 Fig. For the larger sample size of 300,000, it is worth noting that the other four variations of the empirical Bayes method do not seem to perform well compared to other methods, with a positive value for change in RMSE noted in some cases. When the proportion of effect SNPs is 0.001, 'EB-gam-nb' is not behaving as well as seen previously.

In addition to the above, summary statistics were simulated for a quantitative trait in which the effect sizes followed bimodal or skewed distributions and for a binary trait with a normal effect size distribution. For these supplementary investigations, in order to reduce computational burden, the assumption of independent SNPs was maintained, the number of repetitions of each simulation scenario was reduced to 50 and our suggested variations of the empirical Bayes method were excluded from evaluation. The results from assessing the methods using estimated change in RMSE of significant SNPs can be seen in S6-S11 Figs.

Focusing on the $5 \times 10^{-8}$ threshold, very similar conclusions may be deduced for the three different situations considered. Firstly, the extreme unreliability of the conditional likelihood methods is again evident. Out of the six methods evaluated, the empirical Bayes method is clearly the most consistent at achieving less biased association estimates for significant SNPs. In S6, S8 and S10 Figs, it can be seen to perform in a similar manner to the method which obtains the estimated effect sizes of significant SNPs in the discovery GWAS from an independent replication GWAS with a similar number of samples. Its dominance over the other correction methods is most noticeable in the depiction of results corresponding to a quantitative trait in which the effect sizes of SNPs have a skewed distribution. Across all situations, both FIQT and the bootstrap method tend to behave poorly when the proportion of effect SNPs is 0.001 but show improved performances when this proportion is increased to 0.01. However, under the assumption of a skewed effect size distribution, this improvement by the two methods is no longer observed, often resulting in positive values for the estimated change in RMSE.
