## Supplementary Tables 1-8 for "Review and further developments in statistical corrections for Winner’s Curse in genetic association studies"

**S1 Table. The average number and MSE of significant SNPs at two significance thresholds, 5 × 10^-8^ and 5 × 10^-4^, with proportions that indicate the extent of *Winner’s Curse* bias for each simulation scenario.**

| **Simulation scenario** | **1** | **2** | **3** | **4** | **5** | **6** | **7** | **8** |
| --- | --- | --- | --- | --- | --- | --- | --- | --- |
| **sample size *n*** | 30,000 | 300,000 | 30,000 | 300,000 | 30,000 | 300,000 | 30,000 | 300,000 |
| **heritability *h*^2^** | 0.3 | 0.3 | 0.8 | 0.8 | 0.3 | 0.3 | 0.8 | 0.8 |
| **polygenicity *π*** | 0.01 | 0.01 | 0.01 | 0.01 | 0.001 | 0.001 | 0.001 | 0.001 |
| **Metric** |  |  |  |  |  |  |  |  |
| **No. sig. SNPs**  **(5 × 10^-8^)** | 555 | 33,835 | 3,268 | 103,021 | 2,953 | 29,983 | 10,445 | 47,188 |
| **MSE of sig. SNPs**  **(5 × 10^-8^)** | 0.00146 | 0.000025 | 0.000621 | 0.000019 | 0.000398 | 0.000018 | 0.000231 | 0.000017 |
| **Prop. sig. SNPs with *larger* estimate**  **(5 × 10^-8^)** | 0.999 | 0.7771 | 0.9523 | 0.6684 | 0.8339 | 0.6088 | 0.7091 | 0.5651 |
| **Prop. sig. SNPs *significantly* overestimated**  **(5 × 10^-8^)** | 0.9326 | 0.2138 | 0.5715 | 0.13 | 0.3064 | 0.0998 | 0.1655 | 0.0805 |
| **No. sig. SNPs**  **(5 × 10^-4^)** | 16,709 | 114,020 | 36,120 | 212,293 | 19,751 | 54,428 | 32,654 | 69,354 |
| **MSE of sig. SNPs**  **(5 × 10^-4^)** | 0.000843 | 0.000026 | 0.00051 | 0.00002 | 0.00065 | 0.000032 | 0.000445 | 0.000028 |
| **Prop. sig. SNPs with *larger* estimate**  **(5 × 10^-4^)** | 0.9952 | 0.7699 | 0.9378 | 0.6698 | 0.8873 | 0.6625 | 0.774 | 0.6123 |
| **Prop. sig. SNPs *significantly* overestimated**  **(5 × 10^-4^)** | 0.8403 | 0.2216 | 0.5169 | 0.1392 | 0.5512 | 0.2253 | 0.3559 | 0.1814 |

S1 Table details an initial exploration of the various simulation scenarios. The values provided in the table are averages obtained across 100 simulated sets of summary statistiscs for each scenario. The top portion of the table shows the values of the parameters which define each simulation scenario, i.e. sample size, heritability and polygenicity (proportion of effect SNPs). This table corresponds to simulation settings in which a simple correlation structure has been imposed on the set of SNPs. The number of significant SNPs identified for each scenario at two significance thresholds, 5 × 10^-8^ and 5 × 10^-4^, is provided, as well as the naive MSE of these significant SNPs, as defined by Eq (15) in the main manuscript. The table also contains the proportion of significant SNPs that were seen to have a larger estimated effect size than their true effect size, in terms of absolute value. The final row for each threshold provides the proportion of significant SNPs that have significantly overestimated effect sizes. The estimated effect size of a SNP has been defined as significantly overestimated according to Eq (15) in the main text. This metric gives an indication of the extent of *Winner’s Curse* bias present for each simulation scenario and threshold.

**S2 Table. Estimated change in RMSE of significant SNPs at threshold 5 × 10^-8^** **for each method and simulation setting, with a simple correlation structure imposed on the set of SNPs.**

| **Simulation scenario** | **1** | **2** | **3** | **4** | **5** | **6** | **7** | **8** |
| --- | --- | --- | --- | --- | --- | --- | --- | --- |
| **sample size *n*** | 30,000 | 300,000 | 30,000 | 300,000 | 30,000 | 300,000 | 30,000 | 300,000 |
| **heritability *h*^2^** | 0.3 | 0.3 | 0.8 | 0.8 | 0.3 | 0.3 | 0.8 | 0.8 |
| **polygenicity *π*** | 0.01 | 0.01 | 0.01 | 0.01 | 0.01 | 0.001 | 0.001 | 0.001 |
| **Method** |  |  |  |  |  |  |  |  |
| **CL1** | -0.01275 | 0.00312 | 0.00115 | 0.0029 | 0.00661 | 0.00219 | 0.00885 | 0.00158 |
| **CL2** | -0.01543 | 0.00143 | -0.00538 | 0.00174 | 0.00098 | 0.00141 | 0.00466 | 0.00104 |
| **CL3** | -0.01469 | 0.00216 | -0.00256 | 0.00222 | 0.0034 | 0.00171 | 0.00643 | 0.00125 |
| **EB** | -0.01402 | -0.00072 | -0.00776 | -0.00025 | -0.00341 | -0.0001 | -0.00121 | -0.00003 |
| **EB df=7** | -0.01572 | -0.00061 | -0.00789 | -0.00017 | -0.0031 | 0.00048 | -0.00024 | 0.00024 |
| **EB scam** | -0.0146 | -0.00072 | -0.00787 | -0.00025 | -0.00354 | -0.00007 | -0.00116 | 0.00001 |
| **EB gam-po** | -0.01369 | -0.00067 | -0.00804 | -0.0002 | -0.00363 | 0.00083 | -0.00068 | 0.00113 |
| **EB-gam-nb** | -0.01406 | -0.00071 | -0.008 | -0.00024 | -0.00361 | 0.00003 | -0.00117 | 0.00002 |
| **boot** | -0.01409 | -0.00072 | -0.00786 | -0.00026 | -0.00372 | -0.0001 | -0.00124 | -0.00004 |
| **FIQT** | -0.01278 | -0.00073 | -0.00751 | -0.00025 | -0.00374 | 0.00003 | -0.00088 | 0.00007 |

S2 Table provides values for the estimated change in RMSE of significant SNPs, as defined by Eq (14) in the main manuscript, for each *Winner’s Curse* correction method and simulation scenario. This table corresponds to simulation settings in which a simple correlation structure has been imposed on the set of SNPs. The top portion of the table shows the values of the parameters which define each simulation scenario, i.e. sample size, heritability and polygenicity (proportion of effect SNPs). As described in the main manuscript, 100 sets of summary statistics were simulated for each scenario and the correction methods were applied to each set. Thus, the values shown in the remaining portion of the table are the average estimated change in RMSE of significant SNPs due to method implementation across each of these 100 sets. As it is the change in RMSE that has been computed, it is desirable to obtain a negative change, i.e. the RMSE computed upon application of the correction method is smaller than that of the naïve approach. Thus, positive values in the table have been shaded in grey, indicating poor performing methods. The light green shaded cells highlight the method which, on average, resulted in the greatest reduction in RMSE for each simulated scenario.

**S3 Table. Estimated change in MSE of significant SNPs at threshold 5 × 10^-8^** **for each method and simulation setting, with a simple correlation structure imposed on the set of SNPs.**

| **Simulation scenario** | **1** | **2** | **3** | **4** | **5** | **6** | **7** | **8** |
| --- | --- | --- | --- | --- | --- | --- | --- | --- |
| **sample size *n*** | 30,000 | 300,000 | 30,000 | 300,000 | 30,000 | 300,000 | 30,000 | 300,000 |
| **heritability *h*^2^** | 0.3 | 0.3 | 0.8 | 0.8 | 0.3 | 0.3 | 0.8 | 0.8 |
| **polygenicity *π*** | 0.01 | 0.01 | 0.01 | 0.01 | 0.01 | 0.001 | 0.001 | 0.001 |
| **Method** |  |  |  |  |  |  |  |  |
| **CL1** | -0.0008056 | 0.0000409 | 0.0000574 | 0.0000335 | 0.0003062 | 0.0000231 | 0.0003471 | 0.0000154 |
| **CL2** | -0.0009324 | 0.0000163 | -0.0002357 | 0.000018 | 0.0000396 | 0.0000138 | 0.0001631 | 0.0000096 |
| **CL3** | -0.0008989 | 0.0000262 | -0.0001196 | 0.0000241 | 0.0001463 | 0.0000173 | 0.0002365 | 0.0000118 |
| **EB** | -0.0008664 | -0.0000066 | -0.0003212 | -0.0000021 | -0.000124 | -0.0000008 | -0.0000354 | -0.0000002 |
| **EB df=7** | -0.0009446 | -0.0000057 | -0.0003254 | -0.0000014 | -0.0001136 | 0.0000042 | -0.0000072 | 0.000002 |
| **EB scam** | -0.0008935 | -0.0000067 | -0.0003249 | -0.0000021 | -0.0001284 | -0.0000006 | -0.0000339 | 0.0000001 |
| **EB gam-po** | -0.0008511 | -0.0000063 | -0.0003307 | -0.0000017 | -0.0001313 | 0.0000077 | -0.00002 | 0.0000105 |
| **EB-gam-nb** | -0.0008686 | -0.0000066 | -0.0003293 | -0.000002 | -0.0001306 | 0.0000002 | -0.0000342 | 0.0000002 |
| **boot** | -0.0008706 | -0.0000067 | -0.0003247 | -0.0000021 | -0.0001343 | -0.0000008 | -0.0000362 | -0.0000003 |
| **FIQT** | -0.0008063 | -0.0000068 | -0.0003129 | -0.0000021 | -0.0001348 | 0.0000003 | -0.000026 | 0.0000005 |

S3 Table provides values for the estimated change in MSE of significant SNPs, as defined by Eq (14) in the main manuscript, for each *Winner’s Curse* correction method and simulation scenario. This table corresponds to simulation settings in which a simple correlation structure has been imposed on the set of SNPs. The top portion of the table shows the values of the parameters which define each simulation scenario, i.e. sample size, heritability and polygenicity (proportion of effect SNPs). As described in the main manuscript, 100 sets of summary statistics were simulated for each scenario and the correction methods were applied to each set. Thus, the values shown in the remaining portion of the table are the average estimated change in MSE of significant SNPs due to method implementation across each of these 100 sets. As it is the change in MSE that has been computed, it is desirable to obtain a negative change, i.e. the MSE computed upon application of the correction method is smaller than that of the naïve approach. Thus, positive values in the table have been shaded in grey, indicating poor performing methods. The light green shaded cells highlight the method which, on average, resulted in the greatest reduction in MSE for each simulated scenario.

**S4 Table. Estimated relative change in MSE of significant SNPs at threshold 5 × 10^-8^** **for each method and simulation setting, with a simple correlation structure imposed on the set of SNPs.**

| **Simulation scenario** | **1** | **2** | **3** | **4** | **5** | **6** | **7** | **8** |  |
| --- | --- | --- | --- | --- | --- | --- | --- | --- | --- |
| **sample size *n*** | 30,000 | 300,000 | 30,000 | 300,000 | 30,000 | 300,000 | 30,000 | 300,000 |  |
| **heritability *h*^2^** | 0.3 | 0.3 | 0.8 | 0.8 | 0.3 | 0.3 | 0.8 | 0.8 |  |
| **polygenicity *π*** | 0.01 | 0.01 | 0.01 | 0.01 | 0.01 | 0.001 | 0.001 | 0.001 |  |
| **Method** |  |  |  |  |  |  |  |  |  |
| **CL1** | -0.5581 | 1.6391 | 0.1008 | 1.8003 | 0.7879 | 1.3182 | 1.5149 | 0.921 | 0.9405 |
| **CL2** | -0.6479 | 0.6539 | -0.3879 | 0.9694 | 0.1071 | 0.785 | 0.7134 | 0.5745 | 0.346 |
| **CL3** | -0.6239 | 1.0508 | -0.1944 | 1.298 | 0.3796 | 0.9866 | 1.0328 | 0.706 | 0.5794 |
| **EB** | -0.6021 | -0.2657 | -0.5315 | -0.1144 | -0.3127 | -0.0481 | -0.1532 | -0.0122 | -0.255 |
| **EB df=7** | -0.6562 | -0.2282 | -0.5384 | -0.077 | -0.2869 | 0.2406 | -0.03 | 0.1212 | -0.1819 |
| **EB scam** | -0.6209 | -0.2669 | -0.5376 | -0.1143 | -0.3239 | -0.0329 | -0.1468 | 0.006 | -0.255 |
| **EB gam-po** | -0.5916 | -0.2507 | -0.5473 | -0.0911 | -0.3311 | 0.438 | -0.0857 | 0.6273 | -0.104 |
| **EB-gam-nb** | -0.6037 | -0.265 | -0.5449 | -0.1096 | -0.3295 | 0.0145 | -0.1481 | 0.0092 | -0.2471 |
| **boot** | -0.6051 | -0.2687 | -0.5374 | -0.1152 | -0.3384 | -0.0474 | -0.1564 | -0.0177 | -0.2608 |
| **FIQT** | -0.5607 | -0.27 | -0.518 | -0.1146 | -0.3388 | 0.0148 | -0.1117 | 0.0327 | -0.2333 |

S4 Table provides values for the estimated relative change in MSE of significant SNPs for each *Winner’s Curse* correction method and simulation scenario. This table corresponds to simulation settings in which a simple correlation structure has been imposed on the set of SNPs. The top portion of the table shows the values of the parameters which define each simulation scenario, i.e. sample size, heritability and polygenicity (proportion of effect SNPs). As described in the main manuscript, 100 sets of summary statistics were simulated for each scenario and the correction methods were applied to each set. Thus, the values shown in the remaining portion of the table are the average estimated relative change in MSE of significant SNPs due to method implementation across each of these 100 sets. As it is the relative change in MSE that has been computed, it is desirable to obtain a negative change, i.e. the MSE computed upon application of the correction method is smaller than that of the naïve approach. Thus, positive values in the table have been shaded in grey, indicating poor performing methods. The light green shaded cells highlight the method which, on average, resulted in the greatest relative reduction in MSE for each simulated scenario. As the final column contains the mean of each row, it shows that the bootstrap method has the greatest average estimated relative reduction in MSE. This value of -0.2608 suggests that on average, the bootstrap method improves the MSE of significant SNPs by ≈26.08%.

**S5 Table. Estimated change in RMSE of significant SNPs at threshold 5 × 10^-4^** **for each method and simulation setting, with a simple correlation structure imposed on the set of SNPs.**

| **Simulation scenario** | **1** | **2** | **3** | **4** | **5** | **6** | **7** | **8** |
| --- | --- | --- | --- | --- | --- | --- | --- | --- |
| **sample size *n*** | 30,000 | 300,000 | 30,000 | 300,000 | 30,000 | 300,000 | 30,000 | 300,000 |
| **heritability *h*^2^** | 0.3 | 0.3 | 0.8 | 0.8 | 0.3 | 0.3 | 0.8 | 0.8 |
| **polygenicity *π*** | 0.01 | 0.01 | 0.01 | 0.01 | 0.01 | 0.001 | 0.001 | 0.001 |
| **Method** |  |  |  |  |  |  |  |  |
| **CL1** | -0.009565 | 0.001145 | -0.002652 | 0.001324 | -0.004908 | -0.00009 | -0.001573 | -0.000147 |
| **CL2** | -0.010104 | 0.000023 | -0.005838 | 0.000508 | -0.006345 | -0.000457 | -0.003298 | -0.000384 |
| **CL3** | -0.010252 | 0.000501 | -0.004626 | 0.000846 | -0.005981 | -0.000343 | -0.002724 | -0.000317 |
| **EB** | -0.012711 | -0.000697 | -0.007196 | -0.000266 | -0.006605 | -0.000538 | -0.003446 | -0.000398 |
| **EB df=7** | -0.01233 | -0.000651 | -0.007007 | -0.0002 | -0.006539 | -0.000178 | -0.003159 | -0.000058 |
| **EB scam** | -0.012816 | -0.0007 | -0.007199 | -0.000265 | -0.006562 | -0.000502 | -0.003496 | -0.000322 |
| **EB gam-po** | -0.012895 | -0.000683 | -0.007168 | -0.000212 | -0.006375 | 0.00002 | -0.003403 | 0.000366 |
| **EB-gam-nb** | -0.012813 | -0.000701 | -0.007174 | -0.000254 | -0.006551 | -0.000347 | -0.003471 | -0.000134 |
| **boot** | -0.01163 | -0.000702 | -0.007018 | -0.000267 | -0.006536 | -0.000537 | -0.003446 | -0.000406 |
| **FIQT** | -0.011317 | -0.000703 | -0.006862 | -0.000265 | -0.006565 | -0.000444 | -0.003294 | -0.0003 |

S5 Table provides values for the estimated change in RMSE of significant SNPs, as defined by Eq (14) in the main manuscript, for each *Winner’s Curse* correction method and simulation scenario, when a significance threshold of 5 × 10^-4^ is used. This table corresponds to simulation settings in which a simple correlation structure has been imposed on the set of SNPs. The top portion of the table shows the values of the parameters which define each simulation scenario, i.e. sample size, heritability and polygenicity (proportion of effect SNPs). As described in the main manuscript, 100 sets of summary statistics were simulated for each scenario and the correction methods were applied to each set. Thus, the values shown in the remaining portion of the table are the average estimated change in RMSE of significant SNPs due to method implementation across each of these 100 sets. As it is the change in RMSE that has been computed, it is desirable to obtain a negative change, i.e. the RMSE computed upon application of the correction method is smaller than that of the naïve approach. Thus, positive values in the table have been shaded in grey, indicating poor performing methods. The light green shaded cells highlight the method which, on average, resulted in the greatest reduction in RMSE for each simulated scenario.

**S6 Table. Estimated change in MSE of significant SNPs at threshold 5 × 10^-4^ for each method and simulation setting, with a simple correlation structure imposed on the set of SNPs.**

| **Simulation scenario** | **1** | **2** | **3** | **4** | **5** | **6** | **7** | **8** |
| --- | --- | --- | --- | --- | --- | --- | --- | --- |
| **sample size *n*** | 30,000 | 300,000 | 30,000 | 300,000 | 30,000 | 300,000 | 30,000 | 300,000 |
| **heritability *h*^2^** | 0.3 | 0.3 | 0.8 | 0.8 | 0.3 | 0.3 | 0.8 | 0.8 |
| **polygenicity *π*** | 0.01 | 0.01 | 0.01 | 0.01 | 0.01 | 0.001 | 0.001 | 0.001 |
| **Method** |  |  |  |  |  |  |  |  |
| **CL1** | -0.0004635 | 0.000013 | -0.0001124 | 0.0000135 | -0.000226 | -0.000001 | -0.0000638 | -0.0000015 |
| **CL2** | -0.0004842 | 0.0000002 | -0.0002289 | 0.0000048 | -0.0002831 | -0.0000049 | -0.0001281 | -0.0000039 |
| **CL3** | -0.0004898 | 0.0000053 | -0.000187 | 0.0000083 | -0.000269 | -0.0000037 | -0.0001073 | -0.0000033 |
| **EB** | -0.000576 | -0.0000066 | -0.0002724 | -0.0000023 | -0.000293 | -0.0000058 | -0.0001333 | -0.0000041 |
| **EB df=7** | -0.0005633 | -0.0000062 | -0.0002665 | -0.0000017 | -0.0002905 | -0.000002 | -0.0001231 | -0.0000006 |
| **EB scam** | -0.0005794 | -0.0000066 | -0.0002725 | -0.0000023 | -0.0002914 | -0.0000054 | -0.0001351 | -0.0000033 |
| **EB gam-po** | -0.0005819 | -0.0000065 | -0.0002715 | -0.0000018 | -0.0002843 | 0.0000002 | -0.0001318 | 0.000004 |
| **EB-gam-nb** | -0.0005793 | -0.0000066 | -0.0002717 | -0.0000022 | -0.000291 | -0.0000038 | -0.0001342 | -0.0000014 |
| **boot** | -0.0005396 | -0.0000067 | -0.0002669 | -0.0000023 | -0.0002904 | -0.0000058 | -0.0001333 | -0.0000041 |
| **FIQT** | -0.0005286 | -0.0000067 | -0.000262 | -0.0000023 | -0.0002915 | -0.0000048 | -0.0001279 | -0.0000031 |

S6 Table provides values for the estimated change in MSE of significant SNPs, as defined by Eq (14) in the main manuscript, for each *Winner’s Curse* correction method and simulation scenario, when a significance threshold of 5 × 10^-4^ is used. This table corresponds to simulation settings in which a simple correlation structure has been imposed on the set of SNPs. The top portion of the table shows the values of the parameters which define each simulation scenario, i.e. sample size, heritability and polygenicity (proportion of effect SNPs). As described in the main manuscript, 100 sets of summary statistics were simulated for each scenario and the correction methods were applied to each set. Thus, the values shown in the remaining portion of the table are the average estimated change in MSE of significant SNPs due to method implementation across each of these 100 sets. As it is the change in MSE that has been computed, it is desirable to obtain a negative change, i.e. the MSE computed upon application of the correction method is smaller than that of the naïve approach. Thus, positive values in the table have been shaded in grey, indicating poor performing methods. The light green shaded cells highlight the method which, on average, resulted in the greatest reduction in MSE for each simulated scenario.

**S7 Table. Estimated relative change in MSE of significant SNPs at threshold 5 × 10^-4^** **for each method and simulation setting, with a simple correlation structure imposed on the set of SNPs.**

| **Simulation scenario** | **1** | **2** | **3** | **4** | **5** | **6** | **7** | **8** |  |
| --- | --- | --- | --- | --- | --- | --- | --- | --- | --- |
| **sample size *n*** | 30,000 | 300,000 | 30,000 | 300,000 | 30,000 | 300,000 | 30,000 | 300,000 |  |
| **heritability *h*^2^** | 0.3 | 0.3 | 0.8 | 0.8 | 0.3 | 0.3 | 0.8 | 0.8 |  |
| **polygenicity *π*** | 0.01 | 0.01 | 0.01 | 0.01 | 0.01 | 0.001 | 0.001 | 0.001 |  |
| **Method** |  |  |  |  |  |  |  |  |  |
| **CL1** | -0.5507 | 0.5011 | -0.2214 | 0.6834 | -0.348 | -0.0314 | -0.1435 | -0.0544 | -0.0206 |
| **CL2** | -0.5753 | 0.0092 | -0.4511 | 0.2416 | -0.436 | -0.1553 | -0.2885 | -0.1392 | -0.2243 |
| **CL3** | -0.5819 | 0.2069 | -0.3685 | 0.4165 | -0.4143 | -0.118 | -0.2417 | -0.1159 | -0.1521 |
| **EB** | -0.6843 | -0.2552 | -0.5369 | -0.1157 | -0.4512 | -0.1822 | -0.3003 | -0.1443 | -0.3338 |
| **EB df=7** | -0.6693 | -0.2394 | -0.5254 | -0.0876 | -0.4473 | -0.0619 | -0.277 | -0.0217 | -0.2912 |
| **EB scam** | -0.6883 | -0.2564 | -0.5371 | -0.1156 | -0.4487 | -0.1705 | -0.3043 | -0.1177 | -0.3298 |
| **EB gam-po** | -0.6914 | -0.2506 | -0.5352 | -0.0927 | -0.4388 | 0.0083 | -0.2969 | 0.1434 | -0.2691 |
| **EB-gam-nb** | -0.6882 | -0.2567 | -0.5355 | -0.1109 | -0.4481 | -0.1194 | -0.3023 | -0.05 | -0.3139 |
| **boot** | -0.6411 | -0.257 | -0.5261 | -0.1162 | -0.4472 | -0.1818 | -0.3003 | -0.1473 | -0.3271 |
| **FIQT** | -0.628 | -0.2574 | -0.5165 | -0.1155 | -0.4489 | -0.1516 | -0.2882 | -0.11 | -0.3145 |

S7 Table provides values for the estimated relative change in MSE of significant SNPs for each *Winner’s Curse* correction method and simulation scenario, when a significance threshold of 5 × 10^-4^ is used. This table corresponds to simulation settings in which a simple correlation structure has been imposed on the set of SNPs. The top portion of the table shows the values of the parameters which define each simulation scenario, i.e. sample size, heritability and polygenicity (proportion of effect SNPs). As described in the main manuscript, 100 sets of summary statistics were simulated for each scenario and the correction methods were applied to each set. Thus, the values shown in the remaining portion of the table are the average estimated relative change in MSE of significant SNPs due to method implementation across each of these 100 sets. As it is the relative change in MSE that has been computed, it is desirable to obtain a negative change, i.e. the MSE computed upon application of the correction method is smaller than that of the naïve approach. Thus, positive values in the table have been shaded in grey, indicating poor performing methods. The light green shaded cells highlight the method which, on average, resulted in the greatest relative reduction in MSE for each simulated scenario. As the final column contains the mean of each row, it shows that the original empirical Bayes method has the greatest average estimated relative reduction in MSE, when a significance threshold of 5 × 10^-4^ is used. This value of -0.3338 suggests that on average, this form of the empirical Bayes method improves the MSE of significant SNPs by ≈33.38%.

**S8 Table. Estimated MSE of significant SNPs at threshold 5 × 10^-4^** **for each method and data set.**

| **GWAS** | **BMI 1** | **BMI 2** | **T2D 1** | **T2D 2** | **Height 1** | **Height 2** |
| --- | --- | --- | --- | --- | --- | --- |
| **naive** | 0.00244 | 0.00289 | 0.08837 | 0.10361 | 0.00288 | 0.00315 |
| **CL1** | 0.00118 | 0.00145 | 0.01576 | 0.02052 | 0.00296 | 0.00296 |
| **CL2** | 0.00078 | 0.00102 | 0.02869 | 0.03669 | 0.00189 | 0.00189 |
| **CL3** | 0.00087 | 0.00113 | 0.02047 | 0.02674 | 0.00229 | 0.00228 |
| **EB** | 0.00074 | 0.00085 | 0.00478 | 0.00851 | 0.00169 | 0.00173 |
| **EB df=7** | 0.00069 | 0.00079 | 0.0038 | 0.00862 | 0.00206 | 0.00189 |
| **EB scam** | 0.00071 | 0.00077 | 0.00327 | 0.00833 | 0.0017 | 0.00168 |
| **EB gam-po** | 0.00071 | 0.00083 | 0.00381 | 0.00732 | 0.00177 | 0.00168 |
| **EB-gam-nb** | 0.00071 | 0.00081 | 0.00371 | 0.00626 | 0.00173 | 0.00172 |
| **boot** | 0.00077 | 0.00088 | 0.01272 | 0.01819 | 0.00168 | 0.00171 |
| **FIQT** | 0.00077 | 0.00087 | 0.00446 | 0.00781 | 0.0017 | 0.00172 |

S8 Table provides values for the estimated MSE of significant SNPs, as defined by Eq (14) in the main manuscript, using a threshold of 5 × 10^-4^, for each *Winner’s Curse* correction method and UK Biobank data set. The first row of values represents the estimated MSE obtained if the unadjusted estimated effect sizes of the discovery GWAS are used and no correction method has been applied. This is followed by rows which are representative of the use of different correction methods, i.e. the conditional likelihood based methods, the empirical Bayes method and its variations, the proposed bootstrap method and FIQT, respectively. As it is desirable to obtain lower estimated MSE values upon application of a method, values which are greater than their corresponding naïve value have been shaded in grey. The light green shaded cells highlight the method which resulted in the lowest estimated MSE value for each data set.
